# Peripheral Ca_V_2.2 channels in skin regulate prolonged heat hypersensitivity during neuroinflammation

**DOI:** 10.1101/2024.07.13.603149

**Authors:** Anne-Mary N Salib, Meredith J Crane, Amanda M Jamieson, Diane Lipscombe

**Affiliations:** Department of Neuroscience & the Carney Institute for Brain Science Brown University, Providence, RI 02912, USA; Department of Molecular Microbiology and Immunology, Brown University, Providence, RI 02912, USA

## Abstract

Neuroinflammation can lead to chronic maladaptive pain affecting millions of people worldwide. Neurotransmitters, cytokines, and ion channels are implicated in neuro-immune cell signaling but their roles in specific behavioral responses are not fully elucidated. Voltage-gated Ca_V_2.2 channel activity in skin controls rapid and transient heat hypersensitivity induced by intradermal capsaicin via IL-1α cytokine signaling. Ca_V_2.2 channels are not, however, involved in mechanical hypersensitivity that developed in the same animal model. Here, we show that Ca_V_2.2 channels are also critical for heat hypersensitivity induced by the intradermal (*id*) Complete Freund’s Adjuvant (CFA) model of chronic neuroinflammation that involves ongoing cytokine signaling for days. Ongoing CFA-induced cytokine signaling cascades in skin lead to pronounced edema, and hypersensitivity to sensory stimuli. Peripheral Ca_V_2.2 channel activity in skin is required for the full development and week-long time course of heat hypersensitivity induced by *id* CFA. Ca_V_2.2 channels, by contrast, are not involved in paw edema and mechanical hypersensitivity. CFA induced increases in cytokines in hind paws including IL-6 which was dependent on Ca_V_2.2 channel activity. Using IL-6 specific neutralizing antibodies, we show that IL-6 contributes to heat hypersensitivity and, neutralizing both IL-1α and IL-6 was even more effective at reducing the magnitude and duration of CFA-induced heat hypersensitivity. Our findings demonstrate a functional link between Ca_V_2.2 channel activity and the release of IL-6 in skin and show that Ca_V_2.2 channels have a privileged role in the induction and maintenance of heat hypersensitivity during chronic forms of neuroinflammation in skin.

**Significance Statement:** Neuroinflammation can lead to chronic maladaptive pain. Neurotransmitters, ion channels, cytokines, and cytokine receptors are implicated in neuron-immune signaling, but their importance in mediating specific behavioral responses are not fully elucidated. We show that the activity of peripheral Ca_V_2.2 calcium ion channels in skin play a unique role in the induction and maintenance of heat hypersensitivity in the CFA model of prolonged neuroinflammation, without accompanying effects on edema and mechanical hypersensitivity. Blocking peripheral Ca_V_2.2 channel activity reduces local cytokine levels in hind paws injected with CFA including IL-6 and neutralizing IL-6 reduces CFA-induced heat hypersensitivity. Our studies define key signaling molecules that act locally in skin to trigger and maintain heat hypersensitivity during chronic neuroinflammation.

## Introduction

Neuroinflammation and peripheral nerve injury can lead to chronic pain, a condition that affects 1 in 5 adults in the US (Kuehn, 2018). Chronic pain is commonly associated with ongoing neuroinflammation a process which may precede a subset of neurodegenerative diseases (Amor et al., 2010; Zhang et al., 2023). The identity of proteins and signaling molecules that trigger and maintain chronic neuroinflammation may also inform the development of therapies with improved specificity and efficacy (Ebersberger, 2018; Hung et al., 2017; McGowan et al., 2009; Sommer et al., 2018).

The responsiveness of nerve endings to sensory stimuli can change rapidly within seconds, as a warning of potentially damaging insults (Jain et al., 2020). However, prolonged, chronic neuroinflammation is maladaptive and associated with a range of conditions including painful diabetic neuropathy, arthritis, and neurodegeneration (Amor et al., 2010; Caxaria et al., 2023; Dinarello, 2011; Fang et al., 2022; Schaible, 2014; Zhang et al., 2023). A critical rise in intracellular calcium in sensory nerve endings initiates the neuroinflammatory cycle in skin following potentially damaging stimuli. Depending on the stimulus intensity and duration, several long lasting pathological changes can develop including hypersensitivity to sensory stimuli, edema, and increases in proinflammatory cytokines (Cook & McCleskey, 2002; Costigan et al., 2009; Costigan & Woolf, 2000; Louis et al., 1989; Scholz & Woolf, 2002). Calcium-dependent release of pro-inflammatory molecules, including neuropeptides and ATP, from activated peripheral nerve endings act on immune cells triggering cytokine release(Chai et al., 2017; Giuliani et al., 2017; Honore et al., 2009; Wang et al., 2021). In chronic forms of neuroinflammation, a positive feedback cycle of cytokine release and downstream signaling can last for days to weeks, adding to the importance of knowing which cytokines are critical for changes in behavioral sensitivity. The targets of cytokines and their receptors are essential for cellular and behavioral changes in sensitivity (Cook & McCleskey, 2002; Costigan et al., 2009; Costigan & Woolf, 2000; Louis et al., 1989; Scholz & Woolf, 2002) and specific inhibitors of cytokine signaling have been shown to interrupt the inflammatory cycle in skin (Cavalli et al., 2021; Ebbinghaus et al., 2012; Ostrowski et al., 2011; Schaible, 2014; Trier et al., 2019).

CaV2.2 channels in sensory neurons are well-known to play a major role in regulating neuronal excitability and dominate in their control of transmitter release from presynaptic termini in spinal cord and from nerve endings in skin (Chai et al., 2017; DuBreuil et al., 2021; Jayamanne et al., 2013; White & Cousins, 1998). The use of Ca_V_2.2^-/-^ KO mouse models and highly specific toxins, including ω-CgTx MVIIA used clinically to treat otherwise intractable pain, have established the general importance of Ca_V_2.2 channels in the induction and maintenance of pain (Altier & Zamponi, 2004; Atanassoff et al., 2000; Bowersox et al., 1996; Jiang et al., 2013; Khanna et al., 2019; McGivern, 2006; Miljanich, 2004; Schroeder et al., 2006; Scott et al., 2002; Snutch, 2005; Staats et al., 2004; Wang et al., 1998; Wang et al., 2000; Wermeling, 2005). A single dose of ω-CgTx MVIIA, applied intradermally together with capsaicin, specifically prevents the rapid development of heat hypersensitivity without affecting cross-sensitization of mechanoreceptors in the same animals (DuBreuil et al., 2021). IL-1ɑ, an early released proinflammatory alarmin, couples Ca_V_2.2 channel activity to increased excitability of *Trpv1* nociceptors (Salib et al., 2024). Interfering with the activity of IL-1α using a neutralizing antibody significantly reduces capsaicin-induced inflammatory hypersensitivity to both heat and mechanical stimuli (Salib et al., 2024).

The intradermal (*id)* Complete Freund Adjuvant (CFA) mouse model of chronic-like inflammatory pain mimics features of inflammatory pain in humans; specifically robust and persistent edema coupled with prolonged hypersensitivity to heat, cold, and mechanical stimuli with a time course lasting > 7 days (Fehrenbacher et al., 2012; Larson et al., 1986; Stein et al., 1988). CFA, a heat-killed Mycobacterium, triggers inflammation through slow release of antigen at the site of injection (Awate et al., 2013). Here we use the *id* CFA hind paw mouse model of inflammation to establish if Caᵥ2.2 channels are necessary for the development of heat and mechanical hypersensitivity (Fehrenbacher et al., 2012; Kanai et al., 2007; Moy et al., 2018; Yu et al., 2008; Zhang et al., 2021).

CFA can stimulate the release of several cytokines in skin, including IL-1α, IL-1β, IL-4, IL-6, LIF, IL- 10, TNF-α, IL-33, IFNγ, CXCL10, CCL2, CCL4, and MDC ((Cook et al., 2018; Ellis & Bennett, 2013; Hung et al., 2017; Summer et al., 2008; Zhang & An, 2007), Supplementary Table 1) which act on cellular targets with some degree of redundancy (Wong et al., 2016; Zhang & An, 2007). Selective inhibition of cytokine action through the use of neutralizing antibodies (Ebersberger, 2018; Raucci et al., 2019; Zeng et al., 2022) or blocking specific cytokine receptors (Andratsch et al., 2009; Dinarello & van der Meer, 2013; Ebersberger, 2018; Malsch et al., 2014; Pinho-Ribeiro et al., 2017; Sommer et al., 1999; Summer et al., 2008; Zhang et al., 2008) have revealed functional specificity in neuroinflammatory models of pain. For example, IL-1α-β, IL-6, TNF-α, and CCL2 all promote leukocyte infiltration that further perpetuates cytokine production (Cunin et al., 2011; Jain et al., 2020; Perner et al., 2020; Tamari et al., 2021; Trier et al., 2019; Woolf et al., 1997). Sensory neurons are targets of cytokines and they express cytokine receptors that can be upregulated in response to injury or insult (Hollo et al., 2017; Jain et al., 2020; Pinho-Ribeiro et al., 2017; Wang et al., 2009). Sensory nociceptors express receptors for proinflammatory cytokines, including IL-1R1, gp130, and TNFR1, receptors for IL-1α-β, IL-6, and TNF-α respectively (Cunha et al., 2005; Ebbinghaus et al., 2012; Fang et al., 2015; Pinho-Ribeiro et al., 2017) which have been implicated in rapid changes in neuronal excitability including characteristic increased sensitivity to sensory stimuli (Barabas & Stucky, 2013;Kanai et al., 2007; Malsch et al., 2014). In cultured sensory neurons, TNF-α and IL-1β have been shown to upregulate TRPV1 expression and neutralization of TNF-α and IL-1β in vivo reduces thermal hyperalgesia (Schaible, 2014), consistent with their involvement in neuro-immune underlying thermal hyperalgesia (Pinho-Ribeiro et al., 2017; Schaible, 2014).

Nonetheless, local neuroimmune signals associated with inflammatory pain and peripheral hypersensitivity in skin are still not fully characterized. Here, we report that voltage-gated Ca_V_2.2 channels play a critical role in the development of heat hypersensitivity in CFA-induced neuroinflammation in skin. Of 20 cytokines screened during CFA-induced chronic neuroinflammation, nine cytokines were locally elevated in hind paws injected with CFA, and five of these were detected using two independent platforms. IL-6 was found to depend on Ca_V_2.2 channel activity based on two independent cytokine platforms, and we link its presence directly to CFA-induced heat hypersensitivity.

## Materials & Methods

All mice used were bred at Brown University, and all protocols and procedures were approved by the Brown University Institutional Animal Care and Use Committee (IACUC). All experiments were performed in accordance with approved IACUC protocols. 3-6 month male and female mice were used in all experiments, unless otherwise specified. Experimenters were blind to animal genotype, experimental condition, and solution injected and were only unblinded post-analysis. The Ca_V_2.2^-/-^ global deletion (Ca_V_2.2^-/-^ KO) mouse strain used in these studies (*Cacna1b*^tm5.1DiLi^; MGI). contains a STOP cassette in frame, in exon 1 of *Cacna1b* and has been described previously (DuBreuil et al., 2021; Salib et al., 2024). Control mice for comparison to the Ca_V_2.2^-/-^ KO strain were either littermate controls (in all immunophenotyping experiments) or wild-type mice which have been bred in-house in parallel with, and from the same genetic background as Ca_V_2.2^-/-^ KO.

### Hind paw interstitial fluid extraction

Mice were anesthetized using isoflurane (2.0-3.5%) and O₂ (0.6-0.8 LPM) administered continuously via nosecone for the entire hind paw fluid extraction process. The plantar surface of the footpad was injected with 20 µl of 100% CFA (Sigma-Aldrich Cat. #F5881) or 20 µl of 100% CFA + 2 µM ⍵-CgTx MVIIA (Alomone labs, Cat. # C-670) on day 0 using a 30-gauge insulin needle in the center of one hind paw. Interstitial fluid was extracted daily for 7 days post-injection from anesthetized mice. Saline lavage was performed by injecting 30 µl of saline in the ipsilateral (injected) paw and the contralateral (uninjected) paw to control for repeated daily lavages. Fluid was pooled for 8 animals for each experimental condition (ipsilateral lavages pooled, contralateral lavages pooled). Fluid was dispensed into a labeled, pre-chilled Eppendorf tube on dry ice and stored at -80 degrees until used for immunoassay analyses.

### Multiplex bead-based immunoassay (LEGENDplex) fluid analyses

The Custom Mouse Inflammation Panel LEGENDplex (BioLegend, LEGENDplex™) protocol was followed according to the manufacturer’s recommendations. Data was acquired using an Attune NxT Flow Cytometer. We initially ran pilot experiments using hind paw fluid samples for all experimental conditions to determine sample dilution factors and to screen for the presence of 20 cytokines using capture beads targeting: GM-CSF, IL-1α, IL-1β, IL-4, IL-6, IL-9, IL-10, IL-12p70, IL-17A, IL-22, IL-23, IL-33, TNF-α, IFN-γ, CCL2, CXCL10, CCL4, CCL5, and LIF. 5 of 20 cytokines were detected using the LEGENDplex platform in fluid from CFA injected hind paws. Following this initial screen, we selected 12 cytokines for further analyses based on our pilot results and published literature on neuroinflammation in skin (see Supplementary Table 1 for references). The Biolegend LEGENDplex Data Analysis Software Suite (Qognit) was used to determine mean fluorescence intensities and to calculate analyte concentrations using concurrently generated standard curves.

### Electrochemiluminescence spot-based immunoassay fluid validation

Following LEGENDplex analyses, remaining samples were assayed on two custom Meso Scale Discovery (MSD) biomarker panels. All samples underwent the same freeze-thaw frequency and duration cycles. Custom MSD multiplex panel were designed using mouse U-PLEX Biomarker Group 1 markers (TNFα, IL-1β, CCL2, CCL4, CXCL-10, IFNγ, IL-33, IL-10, Il-23, MDC) and was run according to manufacturer’s instructions using a 16-fold dilution. A custom U-PLEX assay (IL-1α and IL-6) was developed and validated using the U-PLEX Development Pack and Multi-Assay SECTOR Plates from MSD. Each capture antibody (R-PLEX anti-mouse IL-1α and U-PLEX anti-mouse IL-6) was incubated with a U-PLEX-coupled Linker for 30 min at room temperature (RT). The plates were read on MSD QuickPlex SQ 120MM Analyzer within 10 mins. To test compatibility, 8pt calibration curves were run in the multiplexed assay as well as each validated singleplex assay and the results compared. The non-specific signals on the plate showed no statistically significant difference to the assay background signal indicating that none of the analytes or detection antibodies bind non-specifically.

Raw data was analyzed using MSD Discovery Workbench analysis software utilizing a four parameter logistic model (4 PL) with 1/y2 weighting.

### Deep punch biopsy for immunophenotyping

In anesthetized mice, we injected the ipsilateral paw with CFA and the contralateral paw was uninjected. Mice were killed using an overdose of isoflurane followed by cervical dislocation one day or three days following injection of CFA, and deep punch biopsies of hind paws were collected from both CFA-injected (ipsilateral) and uninjected (contralateral) paws using 3 mm punch biopsy tools (MedBlade, 2 punches/paw). Paw punches were placed in Miltenyi gentleMACS C-tubes containing 5 ml of RPMI-based enzymatic cocktail kept on ice and containing: 5% FBS, 50U DNase I, and 2 mg/ml collagenase IV. Pooled punch biopsies from 4 mice were considered one biological replicate, with 5-7 biological replicates per condition per time point. Pooled samples were weighed to the nearest 0.0001g prior to digestion and were placed on a prewarmed shaker at 37 degrees for 1 hr at 250 RPM. Single-cell suspensions were obtained using automated dissociation on a gentleMACS dissociator. Following dissociation, 2 mM EDTA was added to neutralize enzymatic activity. Samples were pulsed in a pre-chilled centrifuge. Tube contents were filtered through a 70 μm filter placed on a 50 ml conical tube, gentleMACS C-tube was rinsed with 10 ml filtered 1X PBS + 0.1% BSA. Cells were pelleted at 1300 RPM for 10 mins and the supernatant was removed. The pelleted cells were washed with 5 ml of 1X PBS + 0.1% BSA and spun down at 1300 RPM for 5 mins, resuspended in 500 μl 1X PBS + 0.1% BSA, and counted prior to staining.

Cells were incubated for 10 mins on ice with anti-mouse CD16/CD32 antibody (see Table 1) diluted in 1X PBS + 0.1% BSA to block Fc receptors (Andersen et al., 2016). Surface antibodies were diluted in 1X PBS + 0.1% BSA, then added to the cells in the presence of anti-CD16/CD32 antibody and incubated for 20 mins on ice in the dark. Following a wash, cells were incubated with fixable viability dye diluted in 1X PBS for 20 mins on ice in the dark. Cells were washed and then fixed with 2% PFA for 15 mins on ice in the dark. For intracellular staining, cells were permeabilized with 1X Perm/Wash Buffer for 30 mins on ice. Intracellular antibodies were diluted in 1X Perm/Wash Buffer and incubated with the cells for 30 mins on ice. Cells were washed and resuspended in 1X PBS for acquisition on an Attune NxT Flow Cytometer (ThermoFisher). Analysis was performed using FlowJo Software v10.9.0. Fluorescence minus one (FMO) controls were used to set analysis gates.

**Table 1.** Antibodies and reagents used for deep punch biopsy immunophenotyping. A list of cell surface and intracellular cytokine antibodies used for flow cytometry analysis of cells isolated from deep hind paw punch biopsies (Figs. 2 and Supplementary Fig. S2).

| Target | Fluor | Clone | Lot | Manufacturer | Catalog # | Concentration (final, v/v) |
| --- | --- | --- | --- | --- | --- | --- |
| pan-cytokeratin | Alexa Fluor 488 | AE1/AE3 | 2654135 | ThermoFisher | 53-9003-82 | 1:200 |
| IL-6 | PerCP-eFluor710 | MP5-20F3 | 2609025 | ThermoFisher | 46-7061-82 | 1:100 |
| F4/80 | eFluor660 | BM8 | 2403287 | ThermoFisher | 50-4801-82 | 1:100 |
| MHC class II (IA/IE) | APC-eFluor780 | M5/114.15.2 | 2626372 | ThermoFisher | 47-5321-82 | 1:200 |
| Ly6C | V450 | AL-21 | 6175508 | BD Biosciences | 560594 | 1:200 |
| Ly6G | BV605 | 1A8 | B360656, B387251 | BioLegend | 127639 | 1:200 |
| CD45 | BV711 | 30-F11 | B366812 | BioLegend | 103147 | 1:200 |
| IL-1α | PE | ALF-161 | 2326129 | ThermoFisher | 12-7011-81 | 1:100 |
| CD207 | PE/Dazzle594 | 4C7 | B320091, B394451 | BioLegend | 144211 | 1:200 |
| pro-IL-1β | PE-Cy7 | NJTEN3 | 2396754 | ThermoFisher | 25-7114-82 | 1:200 |
| anti-CD16/CD32 | purified | 93 | 101302 | BioLegend | 101302 | 1:200 |
| Fixable Viability Dye | eFluor506 | n/a | 2290921 | ThermoFisher | 65-0866-14 | 1:1000 |
| Superbright Staining Buffer | n/a | n/a | 2633469 | ThermoFisher | SB-4401-42 | 1:200 |
| Staining buffer | 1x PBS + 0.1% BSA (made in-house) |  |  |  |  |  |
| 2% PFA | 4% PFA (made in-house) diluted in 1x PBS |  |  |  |  |  |
| BD Perm/Wash Buffer 10x solution | diluted to 1x in dH2O | n/a | 2028447 | BD Biosciences | 51-2091KZ |  |

### Behavioral assessments

The experimenter was blind to genotype and experimental condition for all behavioral experiments. Hind paw withdrawal responses to radiant heat were elicited from mouse hind paws using the Hargreaves method (Plantar Analgesia Meter IITC Life Science). Mice were placed in Plexiglas boxes on an elevated glass plate and allowed to habituate for 30 mins prior to testing. A radiant heat source was positioned beneath the mice and aimed using low-intensity visible light to the plantar surface of the hind paw. For all trials, laser settings were: Idle intensity at 5% and active intensity at 50% of maximum. Cut off time = 30 s. Trials began once the high-intensity light source was activated and ended once the mouse withdrew, shook, and/or licked their hind paw following stimulation. Immediately upon meeting response criteria, the high-intensity light source was turned off. The response latency was measured to the nearest 0.01 s for each trial using the built-in timer corresponding to the duration of the high-intensity beam. Three trials were conducted on each hind paw for each mouse, with at least 1 min rest between trials (Hargreaves et al., 1988). An average of 3 trials were used for the analysis. N values reported are the number of mice. After baseline measures, mice were anesthetized with isoflurane during all intradermal injections. Daily behavior was assessed at the same time of day, and uninjected paws were also assessed daily as a control for exposure to anesthetics and repeated measures.

Mechanical paw withdrawal responses were elicited by an automated Von Frey Plantar Aesthesiometer (catalog #37550, Ugo Basile). Mice were placed in an elevated Plexiglas box with a wire mesh bottom and were allowed to habituate for 30 min prior to testing. The plantar surface of hind paws was assessed using a steady ramp of force ranging from 0 to 8g for up to 90 sec. The trial is automatically terminated when the filament buckles or the paw is withdrawn, force and reaction time are captured. After baseline measures, mice were anesthetized with isoflurane during all intradermal injections.

### Data quantification and statistical analysis

Statistical analyses were performed Prism (Version 10; GraphPad). All data are presented as the mean ± SE. Post hoc corrections for multiple comparisons were performed when applicable and as indicated in the results section.

### Data Availability Statement

The Ca_V_2.2^-/-^ KO mouse strain (Cacna1b^tm5.1DiLi^) is described in the MGI database are available by request from the lab.

## Results

We used the *id* CFA model of neuroinflammation in mouse hind paws to establish if there is a direct link between Ca_V_2.2 channel activity, the release of specific cytokines involved in neuroimmune signaling, and correlated behavioral changes in sensitivity to heat and mechanical stimuli. We measured the development and maintenance of well-described features of CFA-induced neuroinflammation in skin including increased sensitivity to heat and mechanical stimulation, increased paw edema, increased immune cell infiltration, and increased levels of cytokines localized to the injected paw. To assess the role of peripheral Ca_V_2.2 channels in neuroinflammation in skin, we measured responses to *id* CFA in hind paws and compared WT mice, global Ca_V_2.2 ^-/-^ knockout mice (Ca_V_2.2 ^-/-^ KO), and WT mice co-injected with the highly specific Ca_V_2.2 blocker ⍵-CgTx MVIIA.

### Ca_V_2.2 channel activity is necessary for prolonged CFA-induced heat hypersensitivity

Baseline behavioral responses to heat and mechanical stimuli applied to hind paws were measured on Day 0 prior to *id* CFA and then daily for one week (Fig. 1). WT mice exhibited characteristic increases in sensitivity to heat (Fig. 1a) and mechanical (Fig. 1c) stimuli within 1 day of *id* CFA, as compared to contralateral paw responses. We observed shorter response latencies to radiant heat and lower response thresholds required to elicit paw withdrawal to mechanical stimuli following *id* CFA, which persisted for the 7-day period that we monitored behavior.

**Figure 1.**
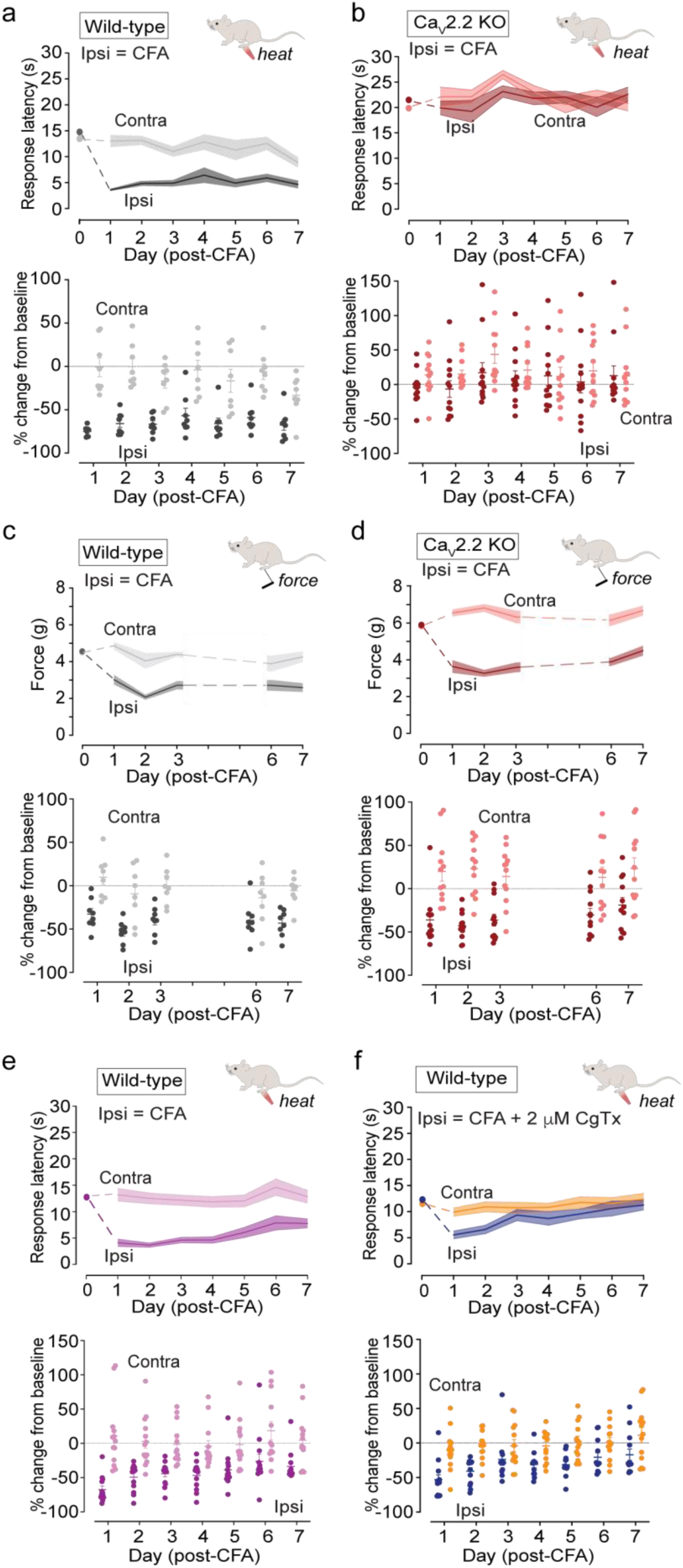
Ca_V_2.2 channel activity is necessary for the development of heat, but not mechanical hypersensitivity triggered by intradermal (*id*) CFA. Daily response latencies to radiant heat **(a, b, e, f)** and withdrawal thresholds to mechanical stimuli **(c, d)** for contra (uninjected) and ipsilateral (CFA injected) hind paws before (Day 0), and for 7 days post *id* 20 μL CFA are shown as mean (solid line; *top*) ± SE (shaded area; *top*) and as individual measurements shown as percentage change from baseline (solid circles; bottom) and mean ± SE (longer and short horizontal lines; *bottom*). Data are shown for two experimental conditions: WT (gray, n = 8, **a,c**) compared to CaV2.2^-/-^ KO (red, n = 12, **b,d**) mice ; and WT CFA (magenta, n = 13, **e**) compared to WT CFA co-injected with 2 μM ⍵-CgTx MVIIA (blue (ipsi), orange (contra), n=13, **f**). **a, b.)** Statistical comparison of % change in withdrawal response latencies to radiant heat for CFA by two-way ANOVA with Tukey HSD correction for multiple comparisons: p(WT ipsi | WT contra) = 0.0001; p(WT ipsi | KO ipsi) = 0.0232; p(KO ipsi | KO contra) = 0.8796. Significance for time | genotype interaction = p < 0.0001. **c, d.)** Same mice used in a, b. Statistical comparison of % change in withdrawal thresholds to mechanical stimuli from baseline following CFA by ANOVA with Tukey HSD correction for multiple comparisons: p(WT ipsi | WT contra) = 0.0073; p(WT ipsi | KO ipsi) = < 0.0001; p(KO ipsi | KO contra) = <0.0001. Significance for time | genotype interaction p = 0.0008. **e-f).** Statistical comparisons of withdrawal latencies to radiant heat following CFA by ANOVA with Tukey HSD correction for multiple comparisons: Time | Injection CFA: p = 0.0100; Time | Injection CFA + CgTx MVIIA: p = 0.2948. Day 3: p(WT CFA ipsi | contra) p = 0.0002, mean = -42.9%; p(WT CFA + CgTx MVIIA ipsi | contra) p = 0.116, mean = -19.9%.

To establish the contribution of Ca_V_2.2 channels to these behavioral responses we assessed the effects of *id* CFA in a Ca_V_2.2 knockout strain (Ca_V_2.2^-/-^ KO) (DuBreuil et al., 2021; Salib et al., 2024) and WT mice injected with *id* ⍵-CgTx MVIIA (Fig. 1). Baseline paw withdrawal response times in Ca_V_2.2^-/-^ KO mice were longer (radiant heat) and occurred at higher thresholds (mechanical) compared to WT (Figs. 1b, 1d). The reduced baseline sensitivity to sensory stimuli in Ca_V_2.2^-/-^ KO mice is expected because synaptic transmission is reduced in Ca_V_2.2 channel-lacking presynaptic nerve endings in the spinal cord (DuBreuil et al., 2021). To normalize for this difference in baseline responses, we also show changes in sensitivity to heat and mechanical stimulation as percentage change from baseline. Within 1 day following *id* CFA, ipsilateral paws of Ca_V_2.2^-/-^ KO mice developed edema (Figs. 2a, 2b) and increased sensitivity to mechanical stimulation based on percentage change from baseline, that were not distinguishable from WT (Figs. 1c, 1d). By comparison, the sensitivity of Ca_V_2.2^-/-^ KO mice to radiant heat did not change following CFA during the 7-day assessment period (p = 0.8796; Fig. 1b). These data, combine with previous studies using the capsaicin model of rapidly developing and transient heat hypersensitivity (DuBreuil et al., 2021; Salib et al., 2024), show that Ca_V_2.2 channels have a privileged and specific role in neuroimmune signaling that triggers long lasting heat hypersensitivity in skin. By contrast, CFA-induced mechanical hypersensitivity and paw edema developed and were maintained by signaling pathways that were independent of Ca_V_2.2 channels.

**Figure 2.**
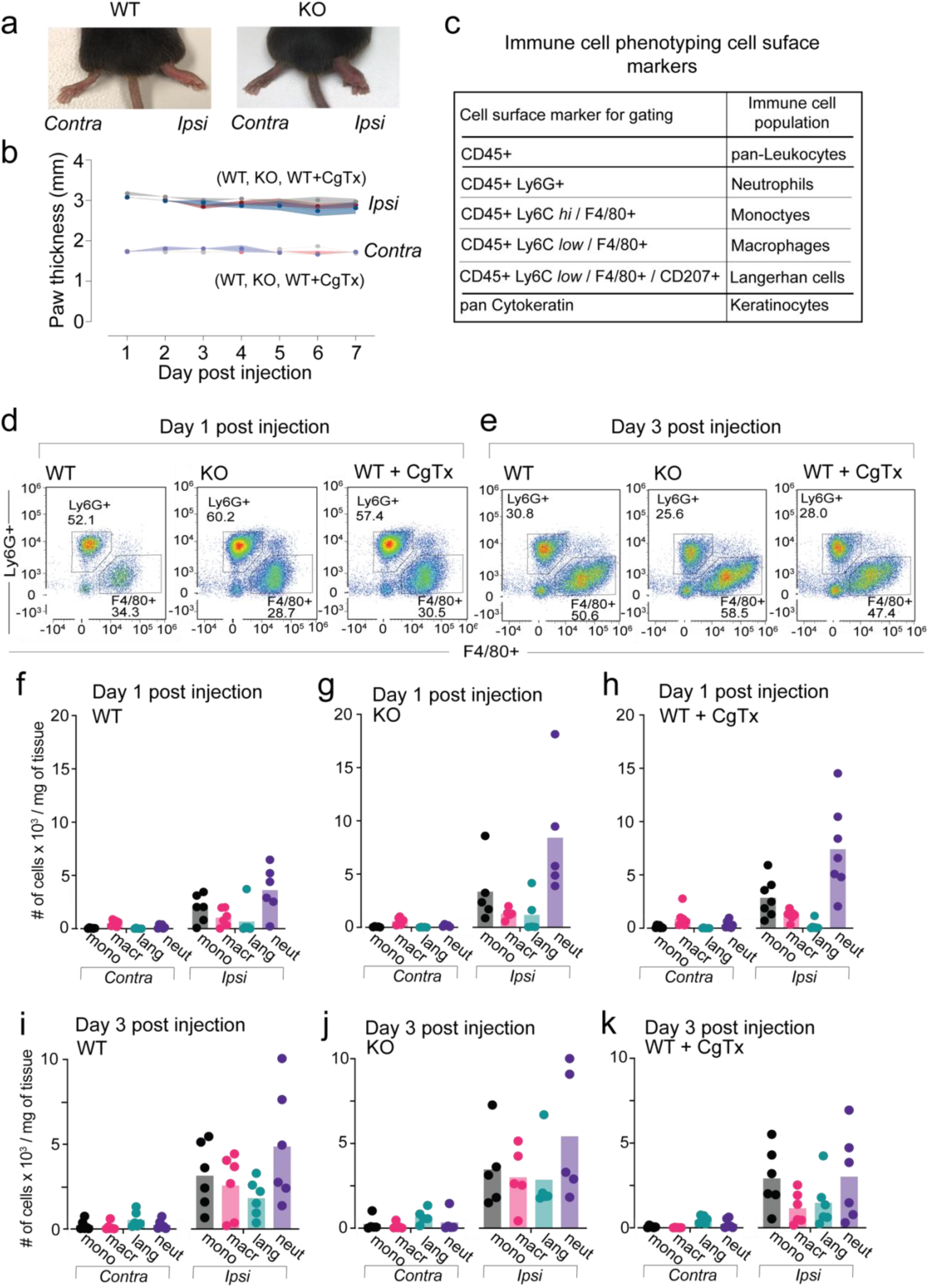
Edema and leukocyte infiltration following intradermal (*id*) CFA is independent of Ca_V_2.2 channel activity. **a.)** Example images of WT and Ca_V_2.2^-/-^ KO mice paws 1 day after CFA injection. **b.)** Paw thickness measured daily using digital calipers. Serial edema measurements of contralateral (control) and ipsilateral (CFA injected) hind paws of mice measured daily for 1 week following 20 μL intradermal *id* injection of CFA in wildtype (WT, gray), CFA in Ca_V_2.2^-/-^ (KO, red), and CFA in WT co- injected with 2 μM ⍵-CgTx MVIIA (WT + CgTx, blue). N=8 mice for each condition. Average values for contralateral paws = 1.74 ± 0.02 mm, and ipsilateral paws = 2.95 ± 0.03 mm. There was no statistical differences in paw thickness of CFA injected paws among the three conditions analysis of variance using two-way ANOVA with for (WT/ KO/ CgTx CFA ipsi paw thickness) | time interaction p = 0.6633. **c.)** Immunophenotyping of leukocytes recruited to the hind paw following CFA injection using a multicolor flow cytometry panel comprising cell surface markers used to identify leukocyte populations (Table 1 and methods). **d-k.)** Two deep punch biopsies were collected from each hind paw, from each animal, 1 day and 3 days post-CFA; samples were pooled (4 pooled ipsilateral paws, 4 contralateral paws per biological replicate; 5-7 biological replicates per condition). Cell counts were normalized to the starting weight of pooled tissue. **d-e.)** Representative flow cytometry plots showing increased neutrophils (Ly6G+) 1 day after CFA injection in the absence of Ca_V_2.2 channel activity when compared to WT controls. Representative flow cytometry plots showing no difference in neutrophils (Ly6G+) 3 days after CFA injection across all 3 experimental conditions. **f-g.)** CD45+ Leukocyte infiltration is increased in ipsilateral (CFA injected) paw tissue on day 1 and day 3 compared to uninjected contralateral paw tissue in WT (WT C=WT contra, WT I= WT ipsi), KO (KO C=KO contra, KO I= KO ipsi), and WT + CgTx MVIIA (WT + Cono C = WT + CgTx contra, WT + Cono I= WT + CgTx ipsi). Analysis of variance by two-way ANOVA with Tukey HSD correction for multiple comparisons **Day 1** : p(WT ipsi | WT contra CD45+) = 0.0414; p(KO ipsi | KO contra CD45+) = 0.0002 p(WT CFA + CgTx MVIIA ipsi | contra CD45+) = 0.0017. **Day 3** : p(WT ipsi | WT contra CD45+) = 0.0006; p(KO ipsi | KO contra CD45+) = 0.0002 p(WT CFA + CgTx MVIIA ipsi | contra CD45+) = 0.0079.Ca_V_2.2 channel activity does not impact leukocyte infiltration into hind paw tissue following *id* CFA on day 1 or day 3; levels of leukocytes in ipsilateral paw tissue are not different when compared across conditions **Day 1** : p(WT ipsi | KO ipsi CD45+) = 0.0665; p(KO ipsi | WT CFA + CgTx MVIIA ipsi CD45+) = 0.5953; p(WT ipsi | WT CFA + CgTx MVIIA ipsi CD45+) = 0.3014. **Day 3** : p(WT ipsi | KO ipsi CD45+) = 0.6876; p(KO ipsi | WT CFA + CgTx MVIIA ipsi CD45+) = 0.1458; p(WT ipsi | WT CFA + CgTx MVIIA ipsi CD45+) = 0.4829. **f.)** Mean cell counts for each cell population 1 day after CFA injection; 1-day following CFA injection Ca_V_2.2 KO and mice co-injected with CgTx MVIIA recruit more neutrophils to the injury site when compared to WT mice injected with CFA alone. Mean neutrophil count normalized to starting weight of the punch Day 1 WT = 3620 ± 529 neutrophils/mg of starting tissue; KO = 8424 ± 1365 neutrophils/mg of starting tissue; WT + CgTx MVIIA = 7414 ± 670 neutrophils/mg of starting tissue; analysis of variance using 2-way ANOVA with Tukey HSD correction for multiple comparisons p(WT CFA ipsi | KO CFA ipsi) = 0.00054; p(WT CFA ipsi | WT CFA + CgTx ipsi) = 0.0191; p(KO CFA ipsi | WT CFA + CgTx ipsi) = 0.7612. **g.)** Mean cell count for each cell population 3 days after CFA injection; no significant difference across cell populations, and specifically neutrophils analysis of variance using 2-way ANOVA with Tukey HSD correction for multiple comparisons on Day 3 p(WT CFA ipsi | KO CFA ipsi) = 0.9080; p(WT CFA ipsi | WT CFA + CgTx ipsi) = 0.3256; p(KO CFA ipsi | WT CFA + CgTx ipsi) = 0.1803.

### Inhibiting peripheral Ca_V_2.2 channels reduces the amplitude and duration of CFA-induced heat hypersensitivity

To assess the importance of Ca_V_2.2 channels in skin, distinct from their role in supporting synaptic transmission at sensory presynaptic sites in the dorsal horn of the spinal cord, we used the highly specific Ca_V_2.2 channel blocker ⍵-CgTx MVIIA (Bowersox et al., 1997; de Souza et al., 2013; Jayamanne et al., 2013; Miljanich, 2004) co-injected together with CFA into hind paws of WT mice (Fig. 1f). Baseline paw withdrawal responses to heat were not different between ipsilateral and contralateral paws (p(ipsi | contra) = 0.5220) (Fig. 1e, 1f). This confirms that *id* ⍵-CgTx MVIIA does not reach the spinal cord and does not interfere with transmission of signals from periphery to central sites that mediate paw withdrawal (DuBreuil et al., 2021). Intradermal ⍵-CgTx MVIIA did, however, reduce both the magnitude and duration of heat hypersensitivity induced by *id* CFA (Fig. 1f). By day 3 ⍵-CgTx MVIIA reduced the effects of *id* CFA significantly compared to control responses (Day 3: WT CFA + CgTx MVIIA ipsi | contra) p = 0.116; Fig. 1f; Day 3: WT CFA ipsi | contra p = 0.0002; Fig. 1e). We also validated the inhibitory effect of ⍵-CgTx MVIIA on CFA-induced heat hypersensitivity in hind paws of WT mice using an independent cohort of outbred C57BL/6 mice (Jackson Labs #000664; Supplementary Fig. S1). These data, combined with previous studies using the *id* capsaicin model of fast, transient heat hypersensitivity (DuBreuil et al., 2021; Salib et al., 2024) show that peripheral Ca_V_2.2 channels contribute selectively to the development of heat hypersensitivity in skin during short and prolonged forms of neuroinflammation. Local inhibition of Ca_V_2.2 channels using a single *id* injection of CgTx MVIIA, coincident with CFA, was sufficient to reduce the magnitude of the early phase, and significantly shorten the time course of heat hypersensitivity in this model of neuroinflammation. By contrast, CFA-induced mechanical hypersensitivity and paw edema developed independent of Ca_V_2.2 channel activity.

### CD45+leukocyte infiltration is independent of Ca_V_2.2 channel activity, but neutrophil density is transiently higher when Ca_V_2.2 channel activity is absent or reduced

We assessed the immune cell composition of hind paw edema in postmortem deep punch biopsies of CFA injected ipsilateral and contralateral hind paws on days 1 and 3 post *id* CFA (Figs. 2c - 2k). We analyzed pooled samples from 4 mice (5 - 7 biological replicates per condition, 20-28 mice per day per condition) and normalized cell counts to the starting weight of pooled tissue (Figs. 2f - 2k; Supplementary Fig. S2 and see methods for flow cytometry gating strategy). Deep punch biopsies of hind paws contained higher levels of CD45+leukocytes 1 and 3 days following *id* CFA as compared to contralateral paws in all three conditions (WT, Ca_V_2.2 ^-/-^, and WT + *id* ⍵-CgTx MVIIA; Figs. 2f - 2k).

CD45+leukocyte infiltration associated with *id* CFA therefore occurs independent of Ca_V_2.2 activity consistent with the importance of CD45+leukocytes in edema (Ghasemlou et al., 2015); a process that is also Ca_V_2.2 channel independent (Figs. 2a, 2b).

Further analysis of CD45+ leukocyte subtypes induced by *id* CFA including monocytes, macrophages, Langerhans cells, and neutrophils (Figs. 2f-2k; also see methods) revealed increased neutrophils in paw punch biopsies of Ca_V_2.2^-/-^ KO and WT mice co-injected ⍵-CgTx MVIIA as compared to WT mice (Figs. 2d, 2f-2h) on day 1: p(WT CFA ipsi | KO CFA ipsi) = 0.00054; p(WT CFA ipsi | WT CFA + CgTx ipsi) = 0.0191; p(KO CFA ipsi | WT CFA + CgTx ipsi) = 0.7612). These data suggest that while not influencing CD45+ leukocyte infiltration overall, Ca_V_2.2 channel activity in skin reduces neutrophil infiltration on day 1 following *id* CFA (Figs. 2f-2h). Cytokines are released by both skin resident immune cells and infiltrating immune cells (Ellis & Bennett, 2013; Ghasemlou et al., 2015; Nguyen &

Soulika, 2019). To further characterize changes in the inflammatory environment that are dependent on Ca_V_2.2 channel activity, we measured cytokine levels in hind paws from pooled lavage fluid extracted daily (Fig. 3).

**Figure 3.**
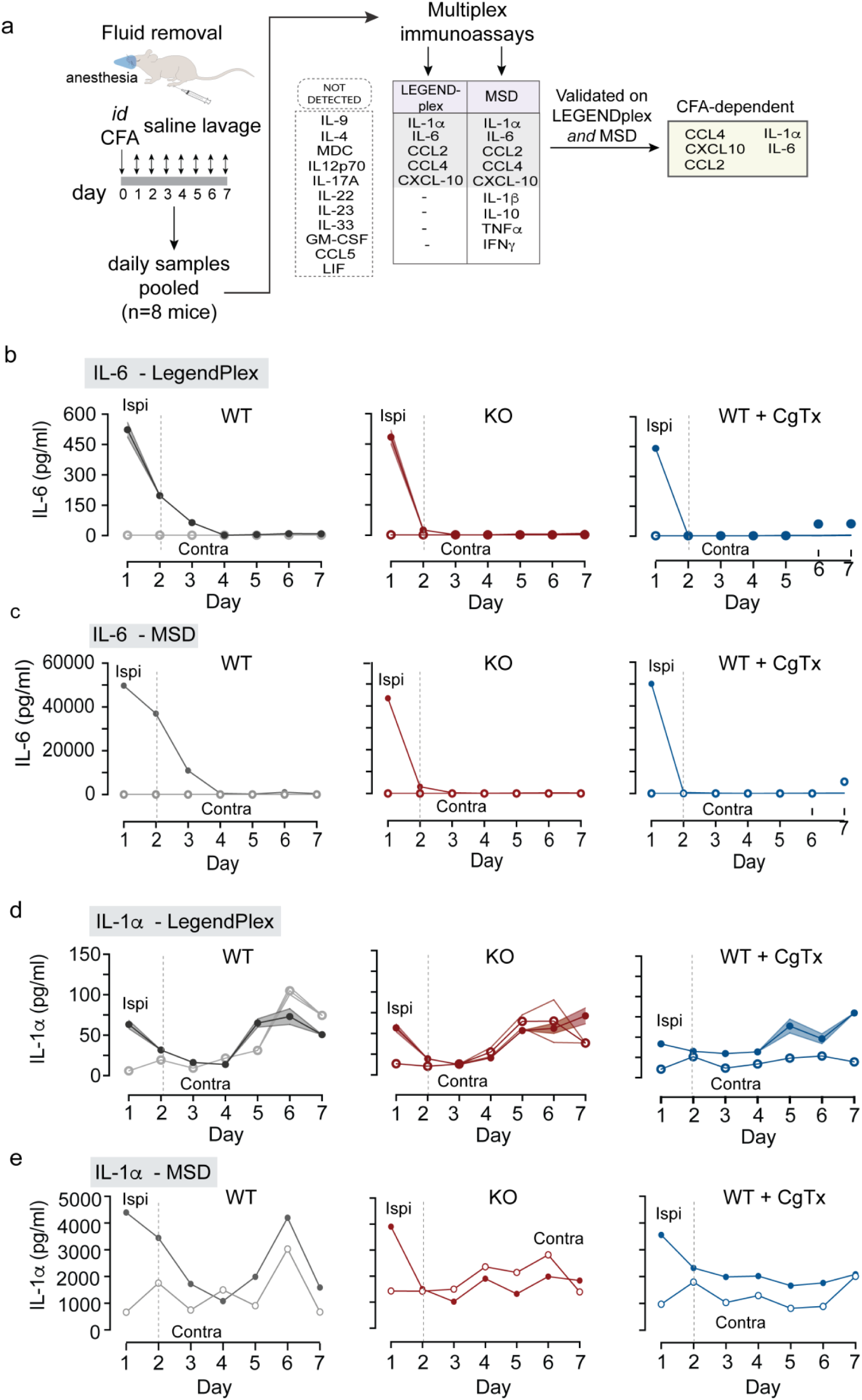
IL-6 levels in CFA treated hind paws are Ca_V_2.2 channel activity-dependent. IL-6 and IL-1α cytokine levels measured daily for 1 week in contralateral and ipsilateral mouse hind paw fluid using customized multiplex LEGENDplex and MSD immunoassays. Ipsilateral paw injected with 20 μL *id* CFA in wildtype (WT, gray), Ca_V_2.2^-/-^ (KO, red), and WT co-injected with 2 μM ⍵-CgTx MVIIA (WT + CgTx, blue). N=8 mice for each condition. **a.)** The experimental protocol summarized: after initial screen for 20 cytokines, 12 cytokines were analyzed in interstitial fluid collected daily from ipsilateral and contralateral hind paws for three experimental conditions. Fluid from 8 mice was pooled for each condition. Five cytokines were CFA-dependent and detected in both assays (IL-6, IL-1α, CCL2, CCL4, and CXCL10, see Supplementary figure S3). **b, d.)** LEGENDplex analyses in technical triplicates of pooled samples from 8 mice per condition. IL-6 (**b.**) and IL-1α (**d.**) levels are elevated following CFA. IL-6 levels in ipsilateral hind paws induced by CFA are Ca_V_2.2 channel dependent. **b.)** Values of IL-6 levels from ipsilateral hind paw fluid are shown as mean ± SE. On day 1: WT = 523 ± 36 pg/ml; Ca_V_2.2^-/-^ KO = 484 ± 34 pg/ml; WT + CgTx = 441 ± 16 pg/ml; Day 2: WT = 197 ± 12 pg/ml; Ca_V_2.2^-/-^ KO = 25 ± 0.93 pg/ml; WT + CgTx = 1.2 ± 0.6 pg/ml; Day 3: WT = 64 ± 15 pg/ml; Ca_V_2.2^-/-^ KO = 1.7 ± 0.1 pg/ml; WT + CgTx MVIIA = 1.17 ± 0.6 pg/ml. Analysis of variance by two-way ANOVA with Tukey HSD correction for multiple comparisons (WT/ KO / CgTx IL-6) | Time interaction p < 0.0001. **Day 2**: p(WT ipsi | KO ipsi IL-6) = 0.0087; p(WT ipsi | WT CFA + CgTx MVIIA ipsi IL-6) = 0.0068. **d.)** Values of IL-1α levels from ipsilateral hind paw fluid are shown as mean ± SE. On day 1 WT = 63 ± 5.0 pg/ml; KO = 56 ± 4.8 pg/ml; WT + CgTx MVIIA = 42 ± 3.1 pg/ml; day 2 WT = 32 ± 1.4 pg/ml; KO = 19 ± 0.2 pg/ml; WT + CgTx MVIIA = 22 ± 11 pg/ml; on day 3 WT = 16 ± 3.8 pg/ml; KO = 12 ± 0.3 pg/ml; WT + CgTx MVIIA = 30 ± 3.5 pg/ml. Analysis of variance by two-way ANOVA with Tukey HSD correction for multiple comparisons (WT/ KO / CgTx IL-1α) | Time interaction p = 0.0001. **Day 2**: p(WT ipsi | KO ipsi IL-1α) = 0.0199; p(WT ipsi | WT CFA + CgTx MVIIA ipsi IL-1α) = 0.6899. **c, e.)** Electrochemiluminescence multiplex spot-based immunoassay (MSD custom R-Plex, U-plex) validation of IL-6 (**c**) and IL-1a (**e**) levels in the same samples used in b & d. Mean values are from 2 technical replicates.

### CFA-dependent increases in proinflammatory cytokines and levels are influenced by Ca_V_2.2 channel activity

Previous studies have shown that Ca_V_2.2 channel activity is necessary to trigger elevated levels of the cytokine IL-1α in the capsaicin-model of fast, reversible neuroinflammation in skin (Salib et al., 2024).

Here we initially screened 20 cytokines in fluid from ipsilateral and contralateral hind paws over a 7- day period and showed 9 cytokines were elevated in hind paws treated with *id* CFA (Fig. 3a). We used this information to select 12 cytokines to generate a custom panel for controlled analysis of fluid extracted from ipsilateral and contralateral hind paws in the *id* CFA model in WT mice, global Ca_V_2.2^-/-^ knockout mice (KO), and WT mice co-injected with the highly specific Ca_V_2.2 blocker ⍵-CgTx MVIIA (Fig. 3; see Supplementary Tables 1 and 2).

Daily fluid samples were analyzed by two independent immunoassay platforms: multiplex bead-based (LEGENDplex; Figs. 3b, 3d; Supplementary Fig. S3) and electrochemiluminescence spot-based (MSD R-plex, Uplex; Figs. 3c, 3e; Supplementary Fig. S3). Using LEGENDplex, we identified elevated levels of 5 of 12 cytokines on at least 1 of 7 days following *id* CFA: IL-1α, IL-6, CXCL10, CCL2, and CCL4 (Fig. 3a; Supplementary Fig. S3). The MSD immunoassay detected elevated levels of the same 5 cytokines detected by the LEGENDplex, in addition to TNF-α, IL-1β, IFNγ, and IL-10. Validating our findings from the LEGENDplex immunoassay on the MSD immunoassay we showed that the MSD immunoassay has greater sensitivity for a subset of cytokines when compared to LEGENDplex (Fig. 3, and Supplementary Fig. S3). The flow cytometry bead-based assay and the electrochemiluminescence-based assay differ in their methods of detection which likely accounts for their different sensitivities. TNF-α, IL-1β, IFNγ, and IL-10 are well-established cytokines involved in CFA induced arthritic models of inflammation in synovial fluid in joints and serum . In mouse hind paws, levels of TNF-α, IL-1β, IFNγ, and IL-10 are likely lower than typically reported where available tissue and volume are more abundant. Ours is one of the few studies we know of that has collected serial measurements of cytokines using pooled hind paw interstitial fluid from mice. We focused our analysis on IL-1α, IL-6, CXCL10, CCL2, and CCL4 which we validated in both immunoassays and of these, the time course of locally elevated IL-6 was consistently longer in WT compared to both Ca_V_2.2^-/-^ KO and *id* ω-CgTx MVIIA conditions (Figs. 3b – 3c).

We therefore focused on IL-6, which was elevated in response to *id* CFA, dependent on Ca_V_2.2 channel activity, detected and validated dynamics in both immunoassay platforms, and for which *in vivo* validated neutralizing antibodies were available. IL-6 levels were elevated on days 1-3 following *id* CFA in WT hind paws. By comparison, in Ca_V_2.2^-/-^ KO and *id* ω-CgTx MVIIA WT mice IL-6 was transiently elevated on day 1 and nearly undetectable on days 2 and 3, day 2: p(WT ipsi | KO ipsi IL- 6) = 0.0087; p(WT ipsi | WT CFA + CgTx MVIIA ipsi IL-6) = 0.0068. (WT| KO| CgTx) | Time interaction p < 0.0001; Figs. 3b - 3c). These data show that IL-6 levels were elevated in hind paw fluid following *id* CFA in two independent immunoassay platforms, and its time course was dependent on peripheral Ca_V_2.2 channel activity.

### IL-6 and IL-1α neutralizing antibodies reduced behavioral responses to id CFA

To establish if there is a link between IL-6 and behavioral responses associated with *id* CFA we used anti-mIL-6-IgG (InvivoGen; Anti-mIL-6-mIgG1e3 InvivoFit InvivoGen; Cat. #mil6-mab15-1) co-injected with CFA (Figs. 4f, 4i) and compared the level of heat hypersensitivity to control animals (CFA alone, Figs. 4e, 4h). Anti-mIL-6-IgG reduced the development of heat hypersensitivity compared to control by 30 - 40% in the first three days following CFA (Figs. 4f, 4i). These data show that IL-6 is released in response to CFA, and that it contributes directly to the development of heat hypersensitivity associated with prolonged neuroinflammation. Anti-mIL-6-IgG did not fully occlude CFA-induced heat hypersensitivity, consistent with the involvement of other cytokines in regulating neuronal responsiveness to sensory stimuli.

**Figure 4.**
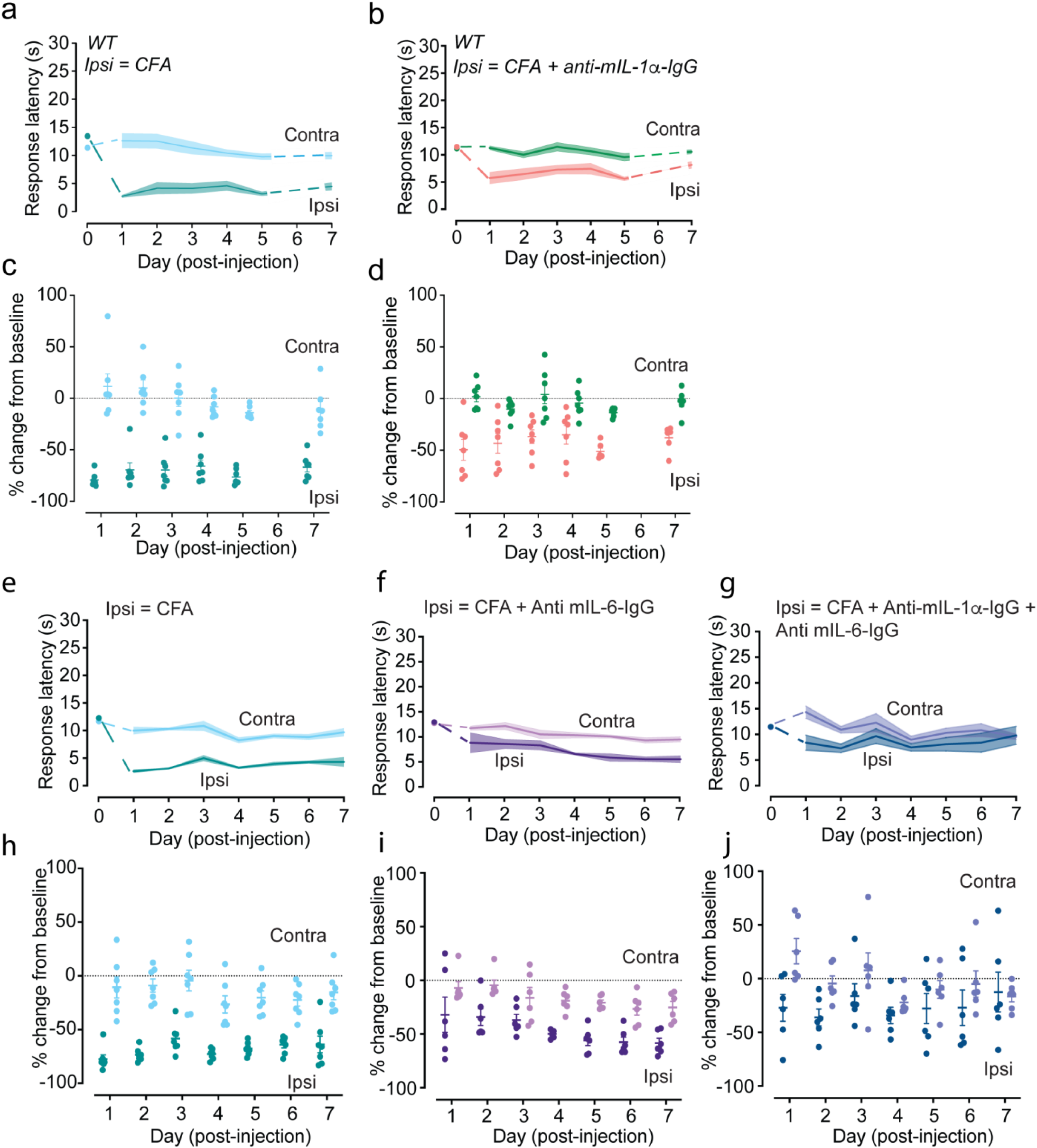
CFA-induced heat hypersensitivity reduced in amplitude and time course by *id* neutralizing antibody to IL-6 and IL-1α. **a-j.)** Withdrawal response latencies (s) to radiant heat in contralateral (contra) and ipsilateral (ipsi) paws shown as mean (lines) ± SE (shaded area) (**a, b, e-g**), and individual responses (solid circles) and mean values (horizontal line) as percent change from baseline (dotted line) (**c, d, h-j**). Measurements obtained immediately prior to (Day 0), and daily for one week after 20 μL *id* CFA together with either: saline (**a, c, e, h**); or 25 μg/ml anti-mIL-1α (b, d); or 100 μg/ml anti-mIL-6 **(f, i**); or 25 μg/ml anti-mIL-1α +100 μg/ml anti-mIL-6 (**g, j**). **a,c.)** control CFA + saline, n=7 (teal); **b,d.)** CFA + 25 μg/ml anti-mIL-1α, n=7 (salmon); Analysis of variance using Two-way ANOVA with Tukey HSD correction for multiple comparisons. interaction between Time | Injection CFA: p =0.0132; Time | Injection CFA + Anti-mIL-1α: p = 0.2504. Day 3 p(WT CFA ipsi | contra) p<0.0001, average percent change from baseline = -69.8%. Day 3 p(WT CFA + Anti-mIL-1α ipsi | contra) p = 0.0036, average percent change from baseline = -40.9%. **e,h.)** control CFA + saline n=7 (teal); **f,i** CFA + 100 μg/ml anti-mIL-6, n = 6 (purple); **g,j.)** CFA + 25 μg/ml anti-mIL-1α +100 μg/ml anti-mIL-6, n = 6 (blue). Analysis of variance using Two-way ANOVA with Tukey HSD correction for multiple comparisons interaction between Time | Injection CFA: p =0.0125; Time | Injection CFA + anti-mIL-6: p = 0.0427; Time | Injection CFA + anti-mIL-1α + anti-mIL-6: p = 0.9181.Day 3 p(WT CFA ipsi | contra) p=0.0008, average percent change from baseline = -58.33%. Day 3 p(WT CFA + anti-mIL-6 ipsi | contra) p =0.0945, average percent change from baseline = -37.00%. Day 3 p(WT CFA + anti-mIL-1α + anti-mIL-6 ipsi | contra) p =0.2613, average percent change from baseline = -13.80%.

IL-1α release occurs within 15 mins of capsaicin-induced stimulation of heat-sensitive nociceptors (Salib et al., 2024) and rapid release in hind paws was Ca_V_2.2 channel activity-dependent (Salib et al., 2024). Further, co-injection of mIL-1ɑ-IgG with capsaicin prevented the development of heat hypersensitivity (Salib et al., 2024). We therefore assessed the effect of anti-mIL-1ɑ-IgG (InvivoGen; Anti-mIL-1α-mIgG1 InvivoFit; Cat. #mil1a-mab9-1) in the CFA model of neuroinflammation and showed a reduction in the degree of heat hypersensitivity (Figs. 4b, 4d). Finally, we showed that the combination of both neutralizing monoclonal antibodies, anti-mIL-1ɑ-IgG and anti-mIL-6-IgG (Fig. 4g), was greater at inhibiting the *id* CFA-associated heat hypersensitivity compared to either anti-mIL-1ɑ- IgG (Fig.4b) or anti-mIL-6-IgG alone (Fig. 4f); by day 3 there was no significant difference in heat hypersensitivity in ipsilateral and contralateral paws (Day 3-Day7) Injection | Time interaction p = 0.5139). These experiments show that both IL-1α and IL-6 contribute to the induction and maintenance of CFA-induced heat hypersensitivity.

When combined with studies on IL-1α which focus on the first 30 mins of acute neuroinflammation following capsaicin exposure (Salib et al., 2024), we conclude that the release of IL-6 and IL-1α are regulated by the activity of peripheral Ca_V_2.2 channels in acute transient and longer-lasting chronic forms of neuroinflammation in skin.

## Discussion

Chronic neuroinflammation in skin can follow peripheral nerve injury, prolonged exposure to damaging stimuli, and is a precursor for a subset of neurodegenerative diseases (Ji et al., 2003; Jiang et al., 2020; Pinho-Ribeiro et al., 2017; Scholz & Woolf, 2007; Zhang & An, 2007). Hallmark behavioral responses of neuroinflammation in skin include long lasting hypersensitivity of sensory neurons to stimuli, including lower response thresholds to noxious heat and mechanical stimuli, as well as perceiving previously innocuous sensory stimuli as painful (Costigan & Woolf, 2000; Ji et al., 2014; Scholz & Woolf, 2007; von Hehn et al., 2012). In addition, edema can develop, and this is associated with increased leukocyte trafficking and extravasation into the injection site (Schober, 2008) as well as increased expression of signaling molecules that perpetuate inflammation triggered by chemokines including CCL2 and CXCL10 (Chen et al., 2019; Ghasemlou et al., 2015; Miller et al., 2009; Prizant et al., 2021; Schober, 2008).

Many molecules including ion channels, neurotransmitters, and cytokines and their respective receptors are implicated in neuroinflammation that causes chronic pain (Supplementary Table 1; (Basbaum et al., 2009; Costigan & Woolf, 2000; Jain et al., 2020; Pinho-Ribeiro et al., 2017; Scholz & Woolf, 2007; Tamari et al., 2021; Trier et al., 2019)). In this study, we identify neuroinflammatory processes and molecules that depend on the activity of voltage-gated Ca_V_2.2 channels. Ca_V_2.2 channels are enriched in*Trpv1* nociceptors where they regulate synaptic transmission in spinal cord (Pitake et al., 2019; White & Cousins, 1998) and in nerve endings skin (DuBreuil et al., 2021). Here, we provide direct evidence that peripheral voltage-gated Ca_V_2.2 channels and locally released cytokines, including IL-6, are key mediators of prolonged heat hypersensitivity that develops in response to neuroinflammation in skin. Our experimental approach was designed to localize the site of action of cytokines and Ca_V_2.2 channels to the same hind paw region where behavioral responses to radiant heat and mechanical stimulation were measured directly. We developed and employed two independent multiplex cytokine assays to confirm the presence of cytokines, including IL-6, in hind paw fluid during the first 3 days after CFA injection, when behavioral changes develop rapidly and are maximal.

### Peripheral Ca_V_2.2 channels – specific role in chronic heat hypersensitivity

Previous studies have shown that peripheral Ca_V_2.2 channels are critical for the development of heat hypersensitivity in skin induced by intradermal capsaicin (DuBreuil et al., 2021); a model of rapid, adaptive increases in sensitivity to sensory stimuli in response to potentially damaging events that peak within 15 mins and reverse within 30 mins (DuBreuil et al., 2021). Here, we show that peripheral Ca_V_2.2 channels have a qualitatively similar role in CFA-induced heat hypersensitivity in skin – which develops rapidly, but with a time course that lasts for days and involves ongoing release of several cytokines (Larson et al., 1986; Pitake et al., 2019; Stein et al., 1988). A single intradermal injection of ⍵-CgTx MVIIA, coincident with CFA injection, was highly effective at reducing the magnitude and shortening the time course of CFA-induced heat hypersensitivity (Figs. 1e, 1f; Supplementary Fig. S1). This suggests that early intervention can be highly effective at reducing the magnitude and time course of the neuroinflammatory response, and that inhibition of peripheral Ca_V_2.2 channels is highly effective at curtailing heat hypersensitivity.

Mice that completely lack Ca_V_2.2 channels fail to develop heat hypersensitivity in response to intradermal CFA (KO; Fig. 1b), but these same mice develop mechanical hypersensitivity (Figs. 1d) and edema (Figs. 2a, 2b) at levels that were indistinguishable from WT. These data demonstrate that the neuroimmune signaling molecules that alter *Trpv1* nociceptor function diverge early in the inflammatory response from those that couple to mechanoreceptors. Our findings are consistent with several behavioral studies that point to the presence of distinct signaling pathways inducing heat and mechanical hypersensitivity (Ebbinghaus et al., 2012; Fuchs et al., 2001; Ghitani et al., 2017; Liu &; Sandkuhler, 2009; Usoskin et al., 2015), although they differ in this regard from White and Cousins (White & Cousins, 1998) who showed that daily injections of ω-CgTx-MVIIA attenuated mechanical hypersensitivity in a peripheral nerve injury model of chronic pain (White & Cousins, 1998).

Mechanoreceptors express voltage-gated calcium channels that are distinct from heat sensitive nociceptors (Cai et al., 2021; Francois et al., 2015; Hoppanova & Lacinova, 2022). In particular, Ca_V_3.2 channels (T-type currents) are expressed at high levels in a class of low-threshold mechanoreceptors (Jung et al., 2023; Sharma et al., 2020; Walcher et al., 2018) and have been implicated in CFA-induced mechanical hypersensitivity (Cai et al., 2021; Picard et al., 2023; Watanabe et al., 2015). Pharmacological and genetic manipulations of Ca_V_3.2 channels reduces CFA-induced mechanical hypersensitivity and reduces CFA-induced edema. Interestingly, levels of IL- 6 were also reduced under conditions of reduced Ca_V_3.2 channel activity (Picard et al., 2023). This suggest that IL-6 signaling can be reduced by targeting Ca_V_3.2 and Ca_V_2.2 calcium ion channels, and is consistent with studies showing that IL-6 has multiple peripheral cellular targets in skin (Brenn et al., 2007; Pinho-Ribeiro et al., 2017; Summer et al., 2008).

The presence of CFA-induced edema in Ca_V_2.2^-/-^ KO mice that failed to develop heat hypersensitivity indicates that CFA is still triggering other neuroinflammatory signaling molecules responsible for edema and mechanical hypersensitivity. Interestingly, CFA-induced edema and heat hypersensitivity were also found to depend differentially on the gp130 protein, a signal transducer that complexes with IL-6Rs (Andratsch et al., 2009). Genetically silencing gp130 in small Na_V_1.8/*SNS*-expressing nociceptors (*SNS-gp130*^−/−^), a population of nociceptors that overlaps substantially with *Trpv1* nociceptors (Andratsch et al., 2009; Jung et al., 2023; Sharma et al., 2020), reduced CFA-induced heat hypersensitivity, while edema remained intact (Andratsch et al., 2009). This report parallels our findings implicating IL-6 signaling specifically in the development of heat hypersensitivity during CFA-induced neuroinflammation (Andratsch et al., 2009).

We also observed an increase of infiltrating leukocytes in CFA-injected paws and this cellular response was independent of Ca_V_2.2 channel activity, consistent with intact edema. However, in hind paws of global Ca_V_2.2^-/-^ KO mice and in WT hind paws treated with ω-CgTx MVIIA, we measured a transient increase in neutrophil infiltration one day after intradermal CFA (Figs. 2d, 2f-h). Additional experiments would be required to establish whether this change is functionally relevant, but a transient upregulation of neutrophils in an inflammatory response has been proposed to protect against the transition from acute to chronic pain in mice (Parisien et al., 2022).This is proposed to reduce the development of sensory neuron hypersensitivity, potentially through secretion of endogenous opioids (Brack et al., 2004; Diatchenko et al., 2022; Parisien et al., 2022; Rittner et al., 2009).

### Cytokines dependent on Ca_V_2.2 channel activity

Sensory nociceptors are the target of a number of proinflammatory cytokines IL-1α/IL-1β, IL-6, and TNF-α which act through their respective receptors IL-1R1, gp130, and TNFR1 respectively (Cunha et al., 2005; Ebbinghaus et al., 2012; Fang et al., 2015; Pinho-Ribeiro et al., 2017). All these cytokine receptors have been implicated in rapid changes in neuronal excitability associated with neuroinflammation including hypersensitivity to sensory stimuli (Barabas & Stucky, 2013; Brenn et al., 2007; Cook et al., 2018; Kanai et al., 2007; Malsch et al., 2014; Tamari et al., 2021; Trier et al., 2019).

We focused on IL-6, which was CFA and Ca_V_2.2 channel dependent, and others have shown its important in the development of acute inflammatory pain (Andratsch et al., 2009; Hurst et al., 2001; Malsch et al., 2014; Murphy et al., 1999; Summer et al., 2008; Wang et al., 2009; Wei et al., 2013) and in modulating the excitability of sensory neurons excitability (Dansereau et al., 2021; Fang et al., 2015; Wang et al., 2009; Wei et al., 2013). Here, we report that the CFA-induced time-course for IL-6 depends on the activity of Ca_V_2.2 channels. Local levels of IL-6 in hind paw fluid induced by CFA were greatly reduced at 2 days, and eliminated 3 days, in hind paws injected with ω−CgTx MVIIA and in Ca_V_2.2^-/-^ KO mice (Figs. 3b - 3c). Our discovery links Ca_V_2.2 channel activation with IL-6 siganling, adding important information about the key neuronal signals that initiate immune cell activation and perpetuate neuroinflammation.

To move beyond correlative observations, and to establish a direct link between cytokine signaling and behavioral hypersensitivity, we used *in vivo* validated neutralizing antibodies to assess behavior affected by local IL-6 signaling. Compared to most studies, which delivered IL-6 neutralizing antibodies or IL-6 receptor antagonists via systemic or intrathecal routes, here we show that local hind paw *id* injection of anti-mIL-6-IgG is highly effective at reducing the degree of CFA-induced heat hypersensitivity. Our results highlight the importance of local cytokine signaling in the development and maintenance of ongoing neuroinflammation. This is consistent with studies of capsaicin-induced heat and mechanical hypersensitivity which were inhibited by intradermal application of a neutralizing antibody to IL-1α (Salib et al., 2024). These findings suggest that local ongoing inflammatory signaling in skin can be interrupted in the periphery, without the need for central or systemic level intervention (DuBreuil et al., 2021; Lee et al., 2019; Salib et al., 2024; White & Cousins, 1998).

Multiple cytokines contribute to ongoing neuroinflammation, but we also show that the combination of neutralizing IL-6 and IL-1α resulted in attenuation of CFA-induced heat hypersensitivity by day 3 (Fig. 4g). While it is very likely that many other cytokines can play a role in CFA-induced neuroinflammation, we find that neutralizing only two of these cytokines is sufficient to significantly shorten the time course of heat hypersensitivity. Cell-surface bound IL-1α acts as an upstream regulator of IL-6 secretion, and depletion of IL-1α using a neutralizing antibody reduces DNA binding activity of NF-kB which stimulates downstream IL-6 transcription (Orjalo et al., 2009). IL-1α is released within the first 15 mins of neuroinflammation induced by intradermal capsaicin (Salib et al., 2024), and it is likely that IL-1α is a key upstream regulator of IL-6 release (Fig. 5).

**Figure 5.**
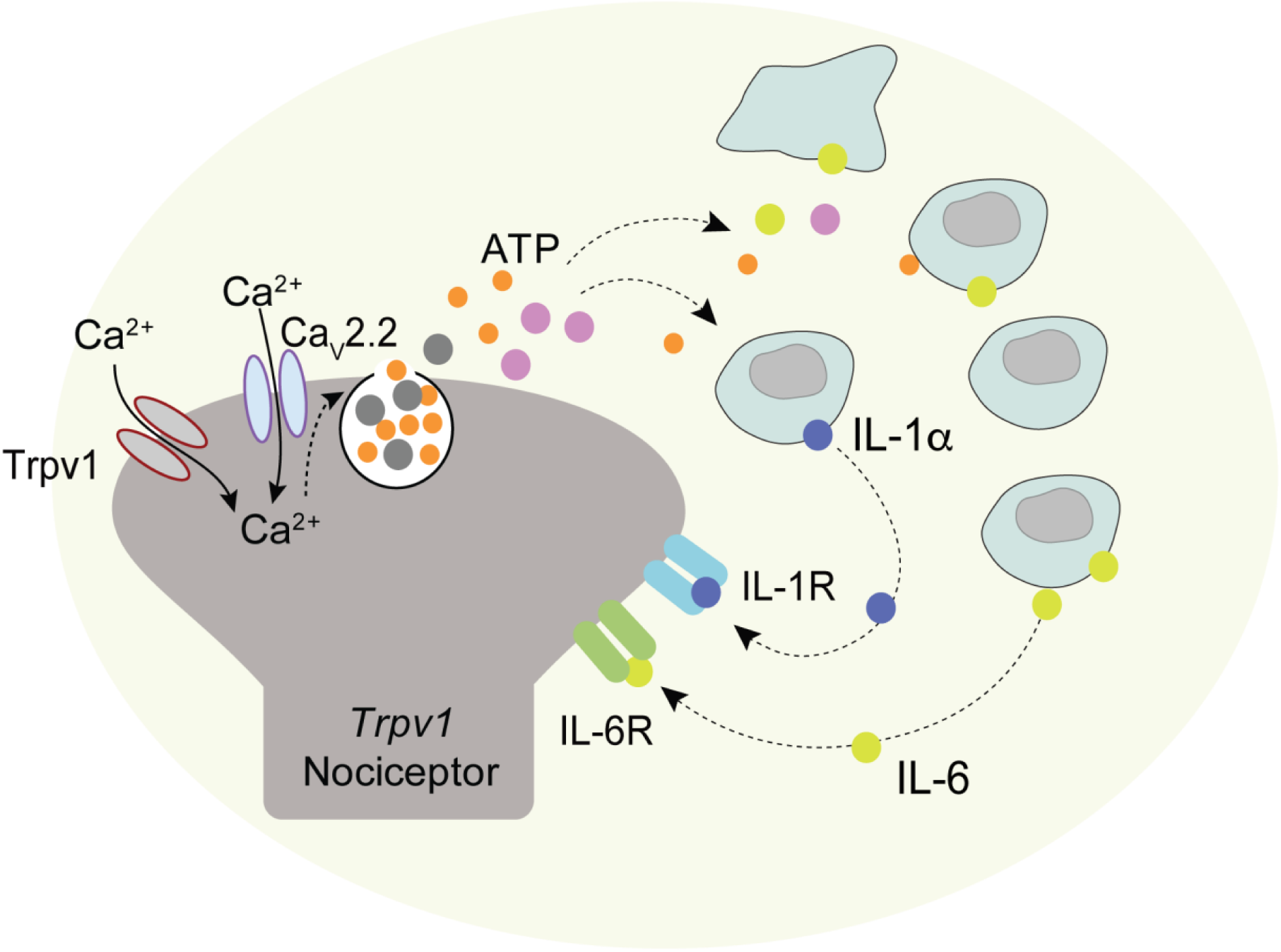
Model focused on the role of IL-6, IL-1α, and Ca_V_2.2 channels in CFA-induced neuroinflammatory signaling. Ca_V_2.2 channels are enriched in *Trpv1* nociceptors including in nerve endings in skin (DuBreuil et al., 2021). *Trpv1* nociceptors depolarize in response to inflammatory agents including CFA. Depolarization results in Ca_V_2.2 channel activation, triggering release of proinflammatory neuropeptides and ATP, and these transmitters can signal to resident and infiltrating immune cell to release cytokines including IL-6 and IL-1α. Increased levels of IL-6 and IL-1α in hind paws act via IL-6R and IL-1R expressed in *Trpv1* nociceptors (Andratsch et al., 2009; Fang et al., 2015; Martin et al., 2021; Schaible, 2014). IL-6 and IL-1α activation of *Trpv1* nociceptors is linked to hypersensitivity to heat associated with both acute and long-lasting neuroinflammation models (Andratsch et al., 2009; Brenn et al., 2007; Fang et al., 2015; Huehnchen et al., 2020; Jeevakumar et al., 2020; Malsch et al., 2014; Wei et al., 2013). Reducing or eliminating Ca_V_2.2 channel activity (Fig. 1), or neutralizing the actions of IL-6 and or IL-1α locally, inhibits the magnitude and time course of CFA-induced heat hypersensitivity (Fig. 4).

Our data presented here add new evidence that links activation of peripheral voltage-gated Ca_V_2.2 channels to local cytokine signaling involved in the development and maintenance of heat hypersensitivity associated with prolonged neuroinflammatory pain. Chronic forms of neuroinflammation involve many signaling molecules that can mediate bidirectional communication between sensory neurons and immune cells. Here, we focus on the importance of peripheral Ca_V_2.2 channels localized to the site of inflammation and their critical role in the induction and maintenance of heat hypersensitivity that lasts for days. We also identify IL-6 as one of likely several cytokines communicating between sensory neurons and immune cells to propagate inflammatory signaling underlying prolonged heat hypersensitivity. Importantly, we show that local inhibition of Ca_V_2.2 channels, or neutralizing the actions of IL-1α and IL-6, at the peripheral origin of neuroinflammation are all highly effective at reducing the amplitude and time course of heat hypersensitivity in skin.

## Supporting information

Salib_Supplementary Figures and Tables 13July2024

## Acknowledgements

Inflammatory cytokines in hind paw fluid samples were validated using the Fluid Biomarkers Laboratory at the Carney Institute for Brain Science at Brown University Center for Alzheimer’s Disease Research. We thank Kristine Pelton for technical assistance.

## REFERENCES

1. Adefegha, S. A., Bottari, N. B., Leal, D. B., de Andrade, C. M., & Schetinger, M. R. (2020). Interferon gamma/interleukin-4 modulation, anti-inflammatory and antioxidant effects of hesperidin in complete Freund’s adjuvant (CFA)-induced arthritis model of rats. Immunopharmacol Immunotoxicol, 42(5), 509–520. 10.1080/08923973.2020.1814806

2. Aguirre, A., Gonzalez-Rodriguez, S., Garcia-Dominguez, M., Lastra, A., Gutierrez-Fernandez, A., Hidalgo, A., Menendez, L., & Baamonde, A. (2020). Dual dose-related effects evoked by CCL4 on thermal nociception after gene delivery or exogenous administration in mice. Biochem Pharmacol, 175, 113903. 10.1016/j.bcp.2020.113903

3. Ahn, D. K., Kim, K. H., Jung, C. Y., Choi, H. S., Lim, E. J., Youn, D. H., & Bae, Y. C. (2005). Role of peripheral group I and II metabotropic glutamate receptors in IL-1beta-induced mechanical allodynia in the orofacial area of conscious rats. Pain, 118(1-2), 53–60. 10.1016/j.pain.2005.07.017

4. Altier, C., & Zamponi, G. W. (2004). Targeting Ca2+ channels to treat pain: T-type versus N-type. Trends Pharmacol Sci, 25(9), 465–470. 10.1016/j.tips.2004.07.004

5. Amor, S., Puentes, F., Baker, D., & van der Valk, P. (2010). Inflammation in neurodegenerative diseases. Immunology, 129(2), 154–169. 10.1111/j.1365-2567.2009.03225.x

6. Andersen, M. N., Al-Karradi, S. N., Kragstrup, T. W., & Hokland, M. (2016). Elimination of erroneous results in flow cytometry caused by antibody binding to Fc receptors on human monocytes and macrophages. Cytometry A, 89(11), 1001–1009. 10.1002/cyto.a.22995

7. Andratsch, M., Mair, N., Constantin, C. E., Scherbakov, N., Benetti, C., Quarta, S., Vogl, C., Sailer, C. A., Uceyler, N., Brockhaus, J., Martini, R., Sommer, C., Zeilhofer, H. U., Muller, W., Kuner, R., Davis, J. B., Rose-John, S., & Kress, M. (2009). A key role for gp130 expressed on peripheral sensory nerves in pathological pain. J Neurosci, 29(43), 13473–13483. 10.1523/JNEUROSCI.1822-09.2009

8. Atanassoff, P. G., Hartmannsgruber, M. W., Thrasher, J., Wermeling, D., Longton, W., Gaeta, R., Singh, T., Mayo, M., McGuire, D., & Luther, R. R. (2000). Ziconotide, a new N-type calcium channel blocker, administered intrathecally for acute postoperative pain. Reg Anesth Pain Med, 25(3), 274–278. 10.1016/s1098-7339(00)90010-5

9. Awate, S., Babiuk, L. A., & Mutwiri, G. (2013). Mechanisms of action of adjuvants. Front Immunol, 4, 114. 10.3389/fimmu.2013.00114

10. Banner, L. R., Patterson, P. H., Allchorne, A., Poole, S., & Woolf, C. J. (1998). Leukemia inhibitory factor is an anti-inflammatory and analgesic cytokine. J Neurosci, 18(14), 5456–5462. 10.1523/JNEUROSCI.18-14-05456.1998

11. Barabas, M. E., & Stucky, C. L. (2013). TRPV1, but not TRPA1, in primary sensory neurons contributes to cutaneous incision-mediated hypersensitivity. Mol Pain, 9, 9. 10.1186/1744-8069-9-9

12. Basbaum, A. I., Bautista, D. M., Scherrer, G., & Julius, D. (2009). Cellular and molecular mechanisms of pain. Cell, 139(2), 267–284. 10.1016/j.cell.2009.09.028

13. Binshtok, A. M., Wang, H., Zimmermann, K., Amaya, F., Vardeh, D., Shi, L., Brenner, G. J., Ji, R. R., Bean, B. P., Woolf, C. J., & Samad, T. A. (2008). Nociceptors are interleukin-1beta sensors. J Neurosci, 28(52), 14062–14073. 10.1523/JNEUROSCI.3795-08.2008

14. Bowersox, S., Mandema, J., Tarczy-Hornoch, K., Miljanich, G., & Luther, R. R. (1997). Pharmacokinetics of SNX-111, a selective N-type calcium channel blocker, in rats and cynomolgus monkeys. Drug Metab Dispos, 25(3), 379–383. https://www.ncbi.nlm.nih.gov/pubmed/9172958

15. Bowersox, S. S., Gadbois, T., Singh, T., Pettus, M., Wang, Y. X., & Luther, R. R. (1996). Selective N-type neuronal voltage-sensitive calcium channel blocker, SNX-111, produces spinal antinociception in rat models of acute, persistent and neuropathic pain. J Pharmacol Exp Ther, 279(3), 1243–1249. https://www.ncbi.nlm.nih.gov/pubmed/8968347

16. Brack, A., Rittner, H. L., Machelska, H., Shaqura, M., Mousa, S. A., Labuz, D., Zollner, C., Schafer, M., & Stein, C. (2004). Endogenous peripheral antinociception in early inflammation is not limited by the number of opioid-containing leukocytes but by opioid receptor expression. Pain, 108(1-2), 67–75. 10.1016/j.pain.2003.12.008

17. Brenn, D., Richter, F., & Schaible, H. G. (2007). Sensitization of unmyelinated sensory fibers of the joint nerve to mechanical stimuli by interleukin-6 in the rat: an inflammatory mechanism of joint pain. Arthritis Rheum, 56(1), 351–359. 10.1002/art.22282

18. Cai, S., Gomez, K., Moutal, A., & Khanna, R. (2021). Targeting T-type/CaV3.2 channels for chronic pain. Transl Res, 234, 20–30. 10.1016/j.trsl.2021.01.002

19. Cavalli, G., Colafrancesco, S., Emmi, G., Imazio, M., Lopalco, G., Maggio, M. C., Sota, J., & Dinarello, C. A. (2021). Interleukin 1alpha: a comprehensive review on the role of IL-1alpha in the pathogenesis and treatment of autoimmune and inflammatory diseases. Autoimmun Rev, 20(3), 102763. 10.1016/j.autrev.2021.102763

20. Caxaria, S., Bharde, S., Fuller, A. M., Evans, R., Thomas, B., Celik, P., Dell’Accio, F., Yona, S., Gilroy, D., Voisin, M. B., Wood, J. N., & Sikandar, S. (2023). Neutrophils infiltrate sensory ganglia and mediate chronic widespread pain in fibromyalgia. Proc Natl Acad Sci U S A, 120(17), e2211631120. 10.1073/pnas.2211631120

21. Chai, Z., Wang, C., Huang, R., Wang, Y., Zhang, X., Wu, Q., Wang, Y., Wu, X., Zheng, L., Zhang, C., Guo, W., Xiong, W., Ding, J., Zhu, F., & Zhou, Z. (2017). CaV2.2 Gates Calcium-Independent but Voltage-Dependent Secretion in Mammalian Sensory Neurons. Neuron, 96(6), 1317–1326 e1314. 10.1016/j.neuron.2017.10.028

22. Chen, Q., Liu, Q., Zhang, Y., Li, S., & Yi, S. (2021). Leukemia inhibitory factor regulates Schwann cell proliferation and migration and affects peripheral nerve regeneration. Cell Death Dis, 12(5), 417. 10.1038/s41419-021-03706-8

23. Chen, Y., Boettger, M. K., Reif, A., Schmitt, A., Uceyler, N., & Sommer, C. (2010). Nitric oxide synthase modulates CFA-induced thermal hyperalgesia through cytokine regulation in mice. Mol Pain, 6, 13. 10.1186/1744-8069-6-13

24. Chen, Y., Yin, D., Fan, B., Zhu, X., Chen, Q., Li, Y., Zhu, S., Lu, R., & Xu, Z. (2019). Chemokine CXCL10/CXCR3 signaling contributes to neuropathic pain in spinal cord and dorsal root ganglia after chronic constriction injury in rats. Neurosci Lett, 694, 20–28. 10.1016/j.neulet.2018.11.021

25. Cook, A. D., Christensen, A. D., Tewari, D., McMahon, S. B., & Hamilton, J. A. (2018). Immune Cytokines and Their Receptors in Inflammatory Pain. Trends Immunol, 39(3), 240–255. 10.1016/j.it.2017.12.003

26. Cook, S. P., & McCleskey, E. W. (2002). Cell damage excites nociceptors through release of cytosolic ATP. Pain, 95(1-2), 41–47. 10.1016/s0304-3959(01)00372-4

27. Costigan, M., Scholz, J., & Woolf, C. J. (2009). Neuropathic pain: a maladaptive response of the nervous system to damage. Annu Rev Neurosci, 32, 1–32. 10.1146/annurev.neuro.051508.135531

28. Costigan, M., & Woolf, C. J. (2000). Pain: molecular mechanisms. J Pain, *1*(3 Suppl), 35-44. http://www.ncbi.nlm.nih.gov/entrez/query.fcgi?cmd=Retrieve&db=PubMed&dopt=Citation&list_uids=14622841

29. Cumberbatch, M., Dearman, R. J., Groves, R. W., Antonopoulos, C., & Kimber, I. (2002). Differential regulation of epidermal langerhans cell migration by interleukins (IL)-1alpha and IL-1beta during irritant- and allergen-induced cutaneous immune responses. Toxicol Appl Pharmacol, 182(2), 126–135. 10.1006/taap.2002.9442

30. Cunha, T. M., Verri, W. A., Jr., Silva, J. S., Poole, S., Cunha, F. Q., & Ferreira, S. H. (2005). A cascade of cytokines mediates mechanical inflammatory hypernociception in mice. Proc Natl Acad Sci U S A, 102(5), 1755–1760. 10.1073/pnas.0409225102

31. Cunin, P., Caillon, A., Corvaisier, M., Garo, E., Scotet, M., Blanchard, S., Delneste, Y., & Jeannin, P. (2011). The tachykinins substance P and hemokinin-1 favor the generation of human memory Th17 cells by inducing IL-1beta, IL-23, and TNF-like 1A expression by monocytes. J Immunol, 186(7), 4175–4182. 10.4049/jimmunol.1002535

32. Dansereau, M. A., Midavaine, E., Begin-Lavallee, V., Belkouch, M., Beaudet, N., Longpre, J. M., Melik-Parsadaniantz, S., & Sarret, P. (2021). Mechanistic insights into the role of the chemokine CCL2/CCR2 axis in dorsal root ganglia to peripheral inflammation and pain hypersensitivity. J Neuroinflammation, 18(1), 79. 10.1186/s12974-021-02125-y

33. de Souza, A. H., Castro, C. J., Jr., Rigo, F. K., de Oliveira, S. M., Gomez, R. S., Diniz, D. M., Borges, M. H., Cordeiro, M. N., Silva, M. A., Ferreira, J., & Gomez, M. V. (2013). An evaluation of the antinociceptive effects of Phalpha1beta, a neurotoxin from the spider Phoneutria nigriventer, and omega-conotoxin MVIIA, a cone snail Conus magus toxin, in rat model of inflammatory and neuropathic pain. Cell Mol Neurobiol, 33(1), 59–67. 10.1007/s10571-012-9871-x

34. DeLeo, J. A., Colburn, R. W., Nichols, M., & Malhotra, A. (1996). Interleukin-6-mediated hyperalgesia/allodynia and increased spinal IL-6 expression in a rat mononeuropathy model. J Interferon Cytokine Res, 16(9), 695–700. 10.1089/jir.1996.16.695

35. Diatchenko, L., Parisien, M., Jahangiri Esfahani, S., & Mogil, J. S. (2022). Omics approaches to discover pathophysiological pathways contributing to human pain. Pain, 163(Suppl 1), S69–S78. 10.1097/j.pain.0000000000002726

36. Dinarello, C. A. (2011). Interleukin-1 in the pathogenesis and treatment of inflammatory diseases. Blood, 117(14), 3720–3732. 10.1182/blood-2010-07-273417

37. Dinarello, C. A., & van der Meer, J. W. (2013). Treating inflammation by blocking interleukin-1 in humans. Semin Immunol, 25(6), 469–484. 10.1016/j.smim.2013.10.008

38. DuBreuil, D. M., Lopez Soto, E. J., Daste, S., Meir, R., Li, D., Wainger, B., Fleischmann, A., & Lipscombe, D. (2021). Heat But Not Mechanical Hypersensitivity Depends on Voltage-Gated CaV2.2 Calcium Channel Activity in Peripheral Axon Terminals Innervating Skin. J Neurosci, 41(36), 7546–7560. 10.1523/JNEUROSCI.0195-21.2021

39. Ebbinghaus, M., Uhlig, B., Richter, F., von Banchet, G. S., Gajda, M., Brauer, R., & Schaible, H. G. (2012). The role of interleukin-1beta in arthritic pain: main involvement in thermal, but not mechanical, hyperalgesia in rat antigen-induced arthritis. Arthritis Rheum, 64(12), 3897–3907. 10.1002/art.34675

40. Ebersberger, A. (2018). The analgesic potential of cytokine neutralization with biologicals. Eur J Pharmacol, 835, 19–30. 10.1016/j.ejphar.2018.07.040

41. Ellis, A., & Bennett, D. L. (2013). Neuroinflammation and the generation of neuropathic pain. Br J Anaesth, 111(1), 26–37. 10.1093/bja/aet128

42. Fang, D., Kong, L. Y., Cai, J., Li, S., Liu, X. D., Han, J. S., & Xing, G. G. (2015). Interleukin-6- mediated functional upregulation of TRPV1 receptors in dorsal root ganglion neurons through the activation of JAK/PI3K signaling pathway: roles in the development of bone cancer pain in a rat model. Pain, 156(6), 1124–1144. 10.1097/j.pain.0000000000000158

43. Fang, X. X., Wang, H., Song, H. L., Wang, J., & Zhang, Z. J. (2022). Neuroinflammation Involved in Diabetes-Related Pain and Itch. Front Pharmacol, 13, 921612. 10.3389/fphar.2022.921612

44. Fehrenbacher, J. C., Vasko, M. R., & Duarte, D. B. (2012). Models of inflammation: Carrageenan- or complete Freund’s Adjuvant (CFA)-induced edema and hypersensitivity in the rat. *Curr Protoc Pharmacol*, Chapter 5, Unit5 4. 10.1002/0471141755.ph0504s56

45. Francois, A., Schuetter, N., Laffray, S., Sanguesa, J., Pizzoccaro, A., Dubel, S., Mantilleri, A., Nargeot, J., Noel, J., Wood, J. N., Moqrich, A., Pongs, O., & Bourinet, E. (2015). The Low-Threshold Calcium Channel Cav3.2 Determines Low-Threshold Mechanoreceptor Function. Cell Rep, 10(3), 370–382. 10.1016/j.celrep.2014.12.042

46. Fuchs, P. N., Meyer, R. A., & Raja, S. N. (2001). Heat, but not mechanical hyperalgesia, following adrenergic injections in normal human skin. Pain, 90(1-2), 15–23. 10.1016/s0304-3959(00)00381-x

47. Ghasemlou, N., Chiu, I. M., Julien, J. P., & Woolf, C. J. (2015). CD11b+Ly6G- myeloid cells mediate mechanical inflammatory pain hypersensitivity. Proc Natl Acad Sci U S A, 112(49), E6808–6817. 10.1073/pnas.1501372112

48. Ghitani, N., Barik, A., Szczot, M., Thompson, J. H., Li, C., Le Pichon, C. E., Krashes, M. J., & Chesler, A. T. (2017). Specialized Mechanosensory Nociceptors Mediating Rapid Responses to Hair Pull. Neuron, 95(4), 944–954 e944. 10.1016/j.neuron.2017.07.024

49. Giuliani, A. L., Sarti, A. C., Falzoni, S., & Di Virgilio, F. (2017). The P2X7 Receptor-Interleukin-1 Liaison. Front Pharmacol, 8, 123. 10.3389/fphar.2017.00123

50. Green, D. P., Limjunyawong, N., Gour, N., Pundir, P., & Dong, X. (2019). A Mast-Cell-Specific Receptor Mediates Neurogenic Inflammation and Pain. Neuron, 101(3), 412–420 e413. 10.1016/j.neuron.2019.01.012

51. Hargreaves, K., Dubner, R., Brown, F., Flores, C., & Joris, J. (1988). A new and sensitive method for measuring thermal nociception in cutaneous hyperalgesia. Pain, 32(1), 77–88. 10.1016/0304-3959(88)90026-7

52. Hollo, K., Ducza, L., Hegyi, Z., Docs, K., Hegedus, K., Bakk, E., Papp, I., Kis, G., Meszar, Z., Bardoczi, Z., & Antal, M. (2017). Interleukin-1 receptor type 1 is overexpressed in neurons but not in glial cells within the rat superficial spinal dorsal horn in complete Freund adjuvant-induced inflammatory pain. J Neuroinflammation, 14(1), 125. 10.1186/s12974-017-0902-x

53. Homma, Y., Brull, S. J., & Zhang, J. M. (2002). A comparison of chronic pain behavior following local application of tumor necrosis factor alpha to the normal and mechanically compressed lumbar ganglia in the rat. Pain, 95(3), 239–246. 10.1016/S0304-3959(01)00404-3

54. Honore, P., Donnelly-Roberts, D., Namovic, M., Zhong, C., Wade, C., Chandran, P., Zhu, C., Carroll, W., Perez-Medrano, A., Iwakura, Y., & Jarvis, M. F. (2009). The antihyperalgesic activity of a selective P2X7 receptor antagonist, A-839977, is lost in IL-1alphabeta knockout mice. Behav Brain Res, 204(1), 77–81. 10.1016/j.bbr.2009.05.018

55. Honore, P., Wade, C. L., Zhong, C., Harris, R. R., Wu, C., Ghayur, T., Iwakura, Y., Decker, M. W., Faltynek, C., Sullivan, J., & Jarvis, M. F. (2006). Interleukin-1alphabeta gene-deficient mice show reduced nociceptive sensitivity in models of inflammatory and neuropathic pain but not post-operative pain. Behav Brain Res, 167(2), 355–364. 10.1016/j.bbr.2005.09.024

56. Hoppanova, L., & Lacinova, L. (2022). Voltage-dependent Ca(V)3.2 and Ca(V)2.2 channels in nociceptive pathways. Pflugers Arch, 474(4), 421–434. 10.1007/s00424-022-02666-y

57. Hua, X., Ge, S., Zhang, M., Mo, F., Zhang, L., Zhang, J., Yang, C., Tai, S., Chen, X., Zhang, L., & Liang, C. (2021). Pathogenic Roles of CXCL10 in Experimental Autoimmune Prostatitis by Modulating Macrophage Chemotaxis and Cytokine Secretion. Front Immunol, 12, 706027. 10.3389/fimmu.2021.706027

58. Huang, J., Gadotti, V. M., Zhang, Z., & Zamponi, G. W. (2021). The IL33 receptor ST2 contributes to mechanical hypersensitivity in mice with neuropathic pain. Mol Brain, 14(1), 35. 10.1186/s13041-021-00752-3

59. Huang, J., Gandini, M. A., Chen, L., M’Dahoma, S., Stemkowski, P. L., Chung, H., Muruve, D. A., & Zamponi, G. W. (2020). Hyperactivity of Innate Immunity Triggers Pain via TLR2-IL-33- Mediated Neuroimmune Crosstalk. Cell Rep, 33(1), 108233. 10.1016/j.celrep.2020.108233

60. Huehnchen, P., Muenzfeld, H., Boehmerle, W., & Endres, M. (2020). Blockade of IL-6 signaling prevents paclitaxel-induced neuropathy in C57Bl/6 mice. Cell Death Dis, 11(1), 45. 10.1038/s41419-020-2239-0

61. Hung, A. L., Lim, M., & Doshi, T. L. (2017). Targeting cytokines for treatment of neuropathic pain. Scand J Pain, 17, 287–293. 10.1016/j.sjpain.2017.08.002

62. Huntula, S., Saegusa, H., Wang, X., Zong, S., & Tanabe, T. (2019). Involvement of N-type Ca(2+) channel in microglial activation and its implications to aging-induced exaggerated cytokine response. Cell Calcium, 82, 102059. 10.1016/j.ceca.2019.102059

63. Hurst, S. M., Wilkinson, T. S., McLoughlin, R. M., Jones, S., Horiuchi, S., Yamamoto, N., Rose-John, S., Fuller, G. M., Topley, N., & Jones, S. A. (2001). Il-6 and its soluble receptor orchestrate a temporal switch in the pattern of leukocyte recruitment seen during acute inflammation. Immunity, 14(6), 705–714. 10.1016/s1074-7613(01)00151-0

64. Jain, A., Hakim, S., & Woolf, C. J. (2020). Unraveling the Plastic Peripheral Neuroimmune Interactome. J Immunol, 204(2), 257–263. 10.4049/jimmunol.1900818

65. Jayamanne, A., Jeong, H. J., Schroeder, C. I., Lewis, R. J., Christie, M. J., & Vaughan, C. W. (2013). Spinal actions of omega-conotoxins, CVID, MVIIA and related peptides in a rat neuropathic pain model. Br J Pharmacol, 170(2), 245–254. 10.1111/bph.12251

66. Jeevakumar, V., Al Sardar, A. K., Mohamed, F., Smithhart, C. M., Price, T., & Dussor, G. (2020). IL-6 induced upregulation of T-type Ca(2+) currents and sensitization of DRG nociceptors is attenuated by MNK inhibition. J Neurophysiol, 124(1), 274–283. 10.1152/jn.00188.2020

67. Ji, R. R., Kohno, T., Moore, K. A., & Woolf, C. J. (2003). Central sensitization and LTP: do pain and memory share similar mechanisms? Trends Neurosci, 26(12), 696–705. 10.1016/j.tins.2003.09.017

68. Ji, R. R., Xu, Z. Z., & Gao, Y. J. (2014). Emerging targets in neuroinflammation-driven chronic pain. Nat Rev Drug Discov, 13(7), 533–548. 10.1038/nrd4334

69. Jiang, B. C., Liu, T., & Gao, Y. J. (2020). Chemokines in chronic pain: cellular and molecular mechanisms and therapeutic potential. Pharmacol Ther, 212, 107581. 10.1016/j.pharmthera.2020.107581

70. Jiang, Y. Q., Andrade, A., & Lipscombe, D. (2013). Spinal morphine but not ziconotide or gabapentin analgesia is affected by alternative splicing of voltage-gated calcium channel CaV2.2 pre-mRNA. Mol Pain, 9, 67. 10.1186/1744-8069-9-67

71. Jung, M., Dourado, M., Maksymetz, J., Jacobson, A., Laufer, B. I., Baca, M., Foreman, O., Hackos, D. H., Riol-Blanco, L., & Kaminker, J. S. (2023). Cross-species transcriptomic atlas of dorsal root ganglia reveals species-specific programs for sensory function. Nat Commun, 14(1), 366. 10.1038/s41467-023-36014-0

72. Junttila, I. S. (2018). Tuning the Cytokine Responses: An Update on Interleukin (IL)-4 and IL-13 Receptor Complexes. Front Immunol, 9, 888. 10.3389/fimmu.2018.00888

73. Kanai, Y., Hara, T., Imai, A., & Sakakibara, A. (2007). Differential involvement of TRPV1 receptors at the central and peripheral nerves in CFA-induced mechanical and thermal hyperalgesia. J Pharm Pharmacol, 59(5), 733–738. 10.1211/jpp.59.5.0015

74. Kang, S., Tanaka, T., Narazaki, M., & Kishimoto, T. (2019). Targeting Interleukin-6 Signaling in Clinic. Immunity, 50(4), 1007–1023. 10.1016/j.immuni.2019.03.026

75. Khanna, R., Yu, J., Yang, X., Moutal, A., Chefdeville, A., Gokhale, V., Shuja, Z., Chew, L. A., Bellampalli, S. S., Luo, S., Francois-Moutal, L., Serafini, M. J., Ha, T., Perez-Miller, S., Park, K. D., Patwardhan, A. M., Streicher, J. M., Colecraft, H. M., & Khanna, M. (2019). Targeting the CaValpha-CaVbeta interaction yields an antagonist of the N-type CaV2.2 channel with broad antinociceptive efficacy. Pain, 160(7), 1644–1661. 10.1097/j.pain.0000000000001524

76. Kuehn, B. (2018). Chronic Pain Prevalence. JAMA, 320(16), 1632. 10.1001/jama.2018.16009

77. Larson, A. A., Brown, D. R., el-Atrash, S., & Walser, M. M. (1986). Pain threshold changes in adjuvant-induced inflammation: a possible model of chronic pain in the mouse. Pharmacol Biochem Behav, 24(1), 49–53. 10.1016/0091-3057(86)90043-2

78. Latourte, A., Cherifi, C., Maillet, J., Ea, H. K., Bouaziz, W., Funck-Brentano, T., Cohen-Solal, M., Hay, E., & Richette, P. (2017). Systemic inhibition of IL-6/Stat3 signalling protects against experimental osteoarthritis. Ann Rheum Dis, 76(4), 748–755. 10.1136/annrheumdis-2016-209757

79. Lee, S., Jo, S., Talbot, S., Zhang, H. B., Kotoda, M., Andrews, N. A., Puopolo, M., Liu, P. W., Jacquemont, T., Pascal, M., Heckman, L. M., Jain, A., Lee, J., Woolf, C. J., & Bean, B. P. (2019). Novel charged sodium and calcium channel inhibitor active against neurogenic inflammation. Elife, 8. 10.7554/eLife.48118

80. Lin, Y., Liu, L., Jiang, H., Zhou, J., & Tang, Y. (2017). Inhibition of interleukin-6 function attenuates the central sensitization and pain behavior induced by osteoarthritis. Eur J Pharmacol, 811, 260–267. 10.1016/j.ejphar.2017.06.032

81. Liu, Y., & Ma, Q. (2011). Generation of somatic sensory neuron diversity and implications on sensory coding. Curr Opin Neurobiol, 21(1), 52–60. 10.1016/j.conb.2010.09.003

82. Liu, Y. L., Zhou, L. J., Hu, N. W., Xu, J. T., Wu, C. Y., Zhang, T., Li, Y. Y., & Liu, X. G. (2007). Tumor necrosis factor-alpha induces long-term potentiation of C-fiber evoked field potentials in spinal dorsal horn in rats with nerve injury: the role of NF-kappa B, JNK and p38 MAPK. Neuropharmacology, 52(3), 708–715. 10.1016/j.neuropharm.2006.09.011

83. Lonnemann, G., Endres, S., Van der Meer, J. W., Cannon, J. G., Koch, K. M., & Dinarello, C. A. (1989). Differences in the synthesis and kinetics of release of interleukin 1 alpha, interleukin 1 beta and tumor necrosis factor from human mononuclear cells. Eur J Immunol, 19(9), 1531–1536. 10.1002/eji.1830190903

84. Louis, S. M., Jamieson, A., Russell, N. J., & Dockray, G. J. (1989). The role of substance P and calcitonin gene-related peptide in neurogenic plasma extravasation and vasodilatation in the rat. Neuroscience, 32(3), 581–586. 10.1016/0306-4522(89)90281-9

85. Macleod, T., Berekmeri, A., Bridgewood, C., Stacey, M., McGonagle, D., & Wittmann, M. (2021). The Immunological Impact of IL-1 Family Cytokines on the Epidermal Barrier. Front Immunol, 12, 808012. 10.3389/fimmu.2021.808012

86. Madisen, L., Mao, T., Koch, H., Zhuo, J. M., Berenyi, A., Fujisawa, S., Hsu, Y. W., Garcia, A. J., 3rd, Gu, X., Zanella, S., Kidney, J., Gu, H., Mao, Y., Hooks, B. M., Boyden, E. S., Buzsaki, G., Ramirez, J. M., Jones, A. R., Svoboda, K., Zeng, H. (2012). A toolbox of Cre-dependent optogenetic transgenic mice for light-induced activation and silencing. Nat Neurosci, 15(5), 793–802. 10.1038/nn.3078

87. Malsch, P., Andratsch, M., Vogl, C., Link, A. S., Alzheimer, C., Brierley, S. M., Hughes, P. A., & Kress, M. (2014). Deletion of interleukin-6 signal transducer gp130 in small sensory neurons attenuates mechanonociception and down-regulates TRPA1 expression. J Neurosci, 34(30), 9845–9856. 10.1523/JNEUROSCI.5161-13.2014

88. Martin, P., Goldstein, J. D., Mermoud, L., Diaz-Barreiro, A., & Palmer, G. (2021). IL-1 Family Antagonists in Mouse and Human Skin Inflammation. Front Immunol, 12, 652846. 10.3389/fimmu.2021.652846

89. McGivern, J. G. (2006). Targeting N-type and T-type calcium channels for the treatment of pain. Drug Discov Today, 11(5-6), 245–253. 10.1016/S1359-6446(05)03662-7

90. McGowan, E., Hoyt, S. B., Li, X., Lyons, K. A., & Abbadie, C. (2009). A peripherally acting Na(v)1.7 sodium channel blocker reverses hyperalgesia and allodynia on rat models of inflammatory and neuropathic pain. Anesth Analg, 109(3), 951–958. 10.1213/ane.0b013e3181b01b02

91. McLoughlin, R. M., Hurst, S. M., Nowell, M. A., Harris, D. A., Horiuchi, S., Morgan, L. W., Wilkinson, T. S., Yamamoto, N., Topley, N., & Jones, S. A. (2004). Differential regulation of neutrophil-activating chemokines by IL-6 and its soluble receptor isoforms. J Immunol, 172(9), 5676–5683. 10.4049/jimmunol.172.9.5676

92. Melemedjian, O. K., Tillu, D. V., Moy, J. K., Asiedu, M. N., Mandell, E. K., Ghosh, S., Dussor, G., & Price, T. J. (2014). Local translation and retrograde axonal transport of CREB regulates IL-6- induced nociceptive plasticity. Mol Pain, 10, 45. 10.1186/1744-8069-10-45

93. Menetski, J., Mistry, S., Lu, M., Mudgett, J. S., Ransohoff, R. M., Demartino, J. A., Macintyre, D. E., & Abbadie, C. (2007). Mice overexpressing chemokine ligand 2 (CCL2) in astrocytes display enhanced nociceptive responses. Neuroscience, 149(3), 706–714. 10.1016/j.neuroscience.2007.08.014

94. Miljanich, G. P. (2004). Ziconotide: neuronal calcium channel blocker for treating severe chronic pain. Curr Med Chem, 11(23), 3029–3040. 10.2174/0929867043363884

95. Miller, R. J., Jung, H., Bhangoo, S. K., & White, F. A. (2009). Cytokine and chemokine regulation of sensory neuron function. Handb Exp Pharmacol(194), 417-449. 10.1007/978-3-540-79090-7_12

96. Moy, J. K., Kuhn, J. L., Szabo-Pardi, T. A., Pradhan, G., & Price, T. J. (2018). eIF4E phosphorylation regulates ongoing pain, independently of inflammation, and hyperalgesic priming in the mouse CFA model. Neurobiol Pain, 4, 45–50. 10.1016/j.ynpai.2018.03.001

97. Murphy, P. G., Ramer, M. S., Borthwick, L., Gauldie, J., Richardson, P. M., & Bisby, M. A. (1999). Endogenous interleukin-6 contributes to hypersensitivity to cutaneous stimuli and changes in neuropeptides associated with chronic nerve constriction in mice. Eur J Neurosci, 11(7), 2243–2253. 10.1046/j.1460-9568.1999.00641.x

98. Nguyen, A. V., & Soulika, A. M. (2019). The Dynamics of the Skin’s Immune System. Int J Mol Sci, 20(8). 10.3390/ijms20081811

99. Oh, S. B., Tran, P. B., Gillard, S. E., Hurley, R. W., Hammond, D. L., & Miller, R. J. (2001). Chemokines and glycoprotein120 produce pain hypersensitivity by directly exciting primary nociceptive neurons. J Neurosci, 21(14), 5027–5035. 10.1523/JNEUROSCI.21-14-05027.2001

100. Orjalo, A. V., Bhaumik, D., Gengler, B. K., Scott, G. K., & Campisi, J. (2009). Cell surface-bound IL- 1alpha is an upstream regulator of the senescence-associated IL-6/IL-8 cytokine network. Proc Natl Acad Sci U S A, 106(40), 17031–17036. 10.1073/pnas.0905299106

101. Ostrowski, S. M., Belkadi, A., Loyd, C. M., Diaconu, D., & Ward, N. L. (2011). Cutaneous denervation of psoriasiform mouse skin improves acanthosis and inflammation in a sensory neuropeptide-dependent manner. J Invest Dermatol, 131(7), 1530–1538. 10.1038/jid.2011.60

102. Parisien, M., Lima, L. V., Dagostino, C., El-Hachem, N., Drury, G. L., Grant, A. V., Huising, J., Verma, V., Meloto, C. B., Silva, J. R., Dutra, G. G. S., Markova, T., Dang, H., Tessier, P. A., Slade, G. D., Nackley, A. G., Ghasemlou, N., Mogil, J. S., Allegri, M., & Diatchenko, L. (2022). Acute inflammatory response via neutrophil activation protects against the development of chronic pain. Sci Transl Med, 14(644), eabj9954. 10.1126/scitranslmed.abj9954

103. Perner, C., Flayer, C. H., Zhu, X., Aderhold, P. A., Dewan, Z. N. A., Voisin, T., Camire, R. B., Chow, O. A., Chiu, I. M., & Sokol, C. L. (2020). Substance P Release by Sensory Neurons Triggers Dendritic Cell Migration and Initiates the Type-2 Immune Response to Allergens. Immunity, 53(5), 1063–1077 e1067. 10.1016/j.immuni.2020.10.001

104. Picard, E., Kerckhove, N., Francois, A., Boudieu, L., Billard, E., Carvalho, F. A., Bogard, G., Gosset, P., Bourdier, J., Aissouni, Y., Bourinet, E., Eschalier, A., Daulhac, L., & Mallet, C. (2023). Role of T CD4(+) cells, macrophages, C-low threshold mechanoreceptors and spinal Ca(v) 3.2 channels in inflammation and related pain-like symptoms in murine inflammatory models. Br J Pharmacol, 180(4), 385–400. 10.1111/bph.15956

105. Pinho-Ribeiro, F. A., Verri, W. A., Jr., & Chiu, I. M. (2017). Nociceptor Sensory Neuron-Immune Interactions in Pain and Inflammation. Trends Immunol, 38(1), 5–19. 10.1016/j.it.2016.10.001

106. Pitake, S., Middleton, L. J., Abdus-Saboor, I., & Mishra, S. K. (2019). Inflammation Induced Sensory Nerve Growth and Pain Hypersensitivity Requires the N-Type Calcium Channel Cav2.2. Front Neurosci, 13, 1009. 10.3389/fnins.2019.01009

107. Prizant, H., Patil, N., Negatu, S., Bala, N., McGurk, A., Leddon, S. A., Hughson, A., McRae, T. D., Gao, Y. R., Livingstone, A. M., Groom, J. R., Luster, A. D., & Fowell, D. J. (2021). CXCL10(+) peripheral activation niches couple preferred sites of Th1 entry with optimal APC encounter. Cell Rep, 36(6), 109523. 10.1016/j.celrep.2021.109523

108. Raucci, F., Iqbal, A. J., Saviano, A., Minosi, P., Piccolo, M., Irace, C., Caso, F., Scarpa, R., Pieretti, S., Mascolo, N., & Maione, F. (2019). IL-17A neutralizing antibody regulates monosodium urate crystal-induced gouty inflammation. Pharmacol Res, 147, 104351. 10.1016/j.phrs.2019.104351

109. Rittner, H. L., Hackel, D., Voigt, P., Mousa, S., Stolz, A., Labuz, D., Schafer, M., Schaefer, M., Stein, C., & Brack, A. (2009). Mycobacteria attenuate nociceptive responses by formyl peptide receptor triggered opioid peptide release from neutrophils. PLoS Pathog, 5(4), e1000362. 10.1371/journal.ppat.1000362

110. Safieh-Garabedian, B., Poole, S., Allchorne, A., Winter, J., & Woolf, C. J. (1995). Contribution of interleukin-1 beta to the inflammation-induced increase in nerve growth factor levels and inflammatory hyperalgesia. Br J Pharmacol, 115(7), 1265–1275. 10.1111/j.1476-5381.1995.tb15035.x

111. Salib, A. N., Crane, M. J., Lee, S. H., Wainger, B. J., Jamieson, A. M., & Lipscombe, D. (2024). Interleukin-1alpha links peripheral Ca(V)2.2 channel activation to rapid adaptive increases in heat sensitivity in skin. Sci Rep, 14(1), 9051. 10.1038/s41598-024-59424-6

112. Sandkuhler, J. (2009). Models and mechanisms of hyperalgesia and allodynia. Physiol Rev, 89(2), 707–758. 10.1152/physrev.00025.2008

113. Sandy-Hindmarch, O., Bennett, D. L., Wiberg, A., Furniss, D., Baskozos, G., & Schmid, A. B. (2022). Systemic inflammatory markers in neuropathic pain, nerve injury, and recovery. Pain, 163(3), 526–537. 10.1097/j.pain.0000000000002386

114. Schaible, H. G. (2014). Nociceptive neurons detect cytokines in arthritis. Arthritis Res Ther, 16(5), 470. 10.1186/s13075-014-0470-8

115. Schober, A. (2008). Chemokines in vascular dysfunction and remodeling. Arterioscler Thromb Vasc Biol, 28(11), 1950–1959. 10.1161/ATVBAHA.107.161224

116. Scholz, J., & Woolf, C. J. (2002). Can we conquer pain? Nat Neurosci, 5 Suppl, 1062–1067. http://www.ncbi.nlm.nih.gov/entrez/query.fcgi?cmd=Retrieve&db=PubMed&dopt=Citation&list_uids=12403987

117. Scholz, J., & Woolf, C. J. (2007). The neuropathic pain triad: neurons, immune cells and glia. Nat Neurosci, 10(11), 1361–1368. 10.1038/nn1992

118. Schroeder, C. I., Doering, C. J., Zamponi, G. W., & Lewis, R. J. (2006). N-type calcium channel blockers: novel therapeutics for the treatment of pain. Med Chem, 2(5), 535–543. 10.2174/157340606778250216

119. Scott, D. A., Wright, C. E., & Angus, J. A. (2002). Actions of intrathecal omega-conotoxins CVID, GVIA, MVIIA, and morphine in acute and neuropathic pain in the rat. Eur J Pharmacol, 451(3), 279–286. 10.1016/s0014-2999(02)02247-1

120. Shamash, S., Reichert, F., & Rotshenker, S. (2002). The cytokine network of Wallerian degeneration: tumor necrosis factor-alpha, interleukin-1alpha, and interleukin-1beta. J Neurosci, 22(8), 3052–3060. 10.1523/JNEUROSCI.22-08-03052.2002

121. Sharma, N., Flaherty, K., Lezgiyeva, K., Wagner, D. E., Klein, A. M., & Ginty, D. D. (2020). The emergence of transcriptional identity in somatosensory neurons. Nature, 577(7790), 392–398. 10.1038/s41586-019-1900-1

122. Silva, J. R., Iftinca, M., Fernandes Gomes, F. I., Segal, J. P., Smith, O. M. A., Bannerman, C. A., Silva Mendes, A., Defaye, M., Robinson, M. E. C., Gilron, I., Cunha, T. M., Altier, C., & Ghasemlou, N. (2022). Skin-resident dendritic cells mediate postoperative pain via CCR4 on sensory neurons. Proc Natl Acad Sci U S A, 119(4). 10.1073/pnas.2118238119

123. Snutch, T. P. (2005). Targeting chronic and neuropathic pain: the N-type calcium channel comes of age. NeuroRx, 2(4), 662–670. 10.1602/neurorx.2.4.662

124. Sommer, C., Leinders, M., & Uceyler, N. (2018). Inflammation in the pathophysiology of neuropathic pain. Pain, 159(3), 595–602. 10.1097/j.pain.0000000000001122

125. Sommer, C., Petrausch, S., Lindenlaub, T., & Toyka, K. V. (1999). Neutralizing antibodies to interleukin 1-receptor reduce pain associated behavior in mice with experimental neuropathy. Neurosci Lett, 270(1), 25–28. 10.1016/s0304-3940(99)00450-4

126. Sonekatsu, M., Taniguchi, W., Yamanaka, M., Nishio, N., Tsutsui, S., Yamada, H., Yoshida, M., & Nakatsuka, T. (2016). Interferon-gamma potentiates NMDA receptor signaling in spinal dorsal horn neurons via microglia-neuron interaction. Mol Pain, 12. 10.1177/1744806916644927

127. Staats, P. S., Yearwood, T., Charapata, S. G., Presley, R. W., Wallace, M. S., Byas-Smith, M., Fisher, R., Bryce, D. A., Mangieri, E. A., Luther, R. R., Mayo, M., McGuire, D., & Ellis, D. (2004). Intrathecal ziconotide in the treatment of refractory pain in patients with cancer or AIDS: a randomized controlled trial. JAMA, 291(1), 63–70. 10.1001/jama.291.1.63

128. Stein, C., Millan, M. J., & Herz, A. (1988). Unilateral inflammation of the hindpaw in rats as a model of prolonged noxious stimulation: alterations in behavior and nociceptive thresholds. Pharmacol Biochem Behav, 31(2), 445–451. 10.1016/0091-3057(88)90372-3

129. Stemkowski, P. L., & Smith, P. A. (2012). Long-term IL-1beta exposure causes subpopulation-dependent alterations in rat dorsal root ganglion neuron excitability. J Neurophysiol, 107(6), 1586–1597. 10.1152/jn.00587.2011

130. Summer, G. J., Romero-Sandoval, E. A., Bogen, O., Dina, O. A., Khasar, S. G., & Levine, J. D. (2008). Proinflammatory cytokines mediating burn-injury pain. Pain, 135(1-2), 98–107. 10.1016/j.pain.2007.05.012

131. Tamari, M., Ver Heul, A. M., & Kim, B. S. (2021). Immunosensation: Neuroimmune Cross Talk in the Skin. Annu Rev Immunol, 39, 369–393. 10.1146/annurev-immunol-101719-113805

132. Trier, A. M., Mack, M. R., & Kim, B. S. (2019). The Neuroimmune Axis in Skin Sensation, Inflammation, and Immunity. J Immunol, 202(10), 2829–2835. 10.4049/jimmunol.1801473

133. Usoskin, D., Furlan, A., Islam, S., Abdo, H., Lonnerberg, P., Lou, D., Hjerling-Leffler, J., Haeggstrom, J., Kharchenko, O., Kharchenko, P. V., Linnarsson, S., & Ernfors, P. (2015). Unbiased classification of sensory neuron types by large-scale single-cell RNA sequencing. Nat Neurosci, 18(1), 145–153. 10.1038/nn.3881

134. Van Steenwinckel, J., Auvynet, C., Sapienza, A., Reaux-Le Goazigo, A., Combadiere, C., & Melik Parsadaniantz, S. (2015). Stromal cell-derived CCL2 drives neuropathic pain states through myeloid cell infiltration in injured nerve. Brain Behav Immun, 45, 198–210. 10.1016/j.bbi.2014.10.016

135. von Hehn, C. A., Baron, R., & Woolf, C. J. (2012). Deconstructing the neuropathic pain phenotype to reveal neural mechanisms. Neuron, 73(4), 638–652. 10.1016/j.neuron.2012.02.008

136. Walcher, J., Ojeda-Alonso, J., Haseleu, J., Oosthuizen, M. K., Rowe, A. H., Bennett, N. C., & Lewin, G. R. (2018). Specialized mechanoreceptor systems in rodent glabrous skin. J Physiol, 596(20), 4995–5016. 10.1113/JP276608

137. Wang, A., Shi, X., Yu, R., Qiao, B., Yang, R., & Xu, C. (2021). The P2X(7) Receptor Is Involved in Diabetic Neuropathic Pain Hypersensitivity Mediated by TRPV1 in the Rat Dorsal Root Ganglion. Front Mol Neurosci, 14, 663649. 10.3389/fnmol.2021.663649

138. Wang, X. M., Hamza, M., Wu, T. X., & Dionne, R. A. (2009). Upregulation of IL-6, IL-8 and CCL2 gene expression after acute inflammation: Correlation to clinical pain. Pain, 142(3), 275–283. 10.1016/j.pain.2009.02.001

139. Wang, Y. X., Bezprozvannaya, S., Bowersox, S. S., Nadasdi, L., Miljanich, G., Mezo, G., Silva, D., Tarczy-Hornoch, K., & Luther, R. R. (1998). Peripheral versus central potencies of N-type voltage-sensitive calcium channel blockers. Naunyn Schmiedebergs Arch Pharmacol, 357(2), 159–168. 10.1007/pl00005150

140. Wang, Y. X., Pettus, M., Gao, D., Phillips, C., & Scott Bowersox, S. (2000). Effects of intrathecal administration of ziconotide, a selective neuronal N-type calcium channel blocker, on mechanical allodynia and heat hyperalgesia in a rat model of postoperative pain. Pain, 84(2- 3), 151–158. 10.1016/s0304-3959(99)00197-9

141. Watanabe, M., Ueda, T., Shibata, Y., Kumamoto, N., Shimada, S., & Ugawa, S. (2015). Expression and Regulation of Cav3.2 T-Type Calcium Channels during Inflammatory Hyperalgesia in Mouse Dorsal Root Ganglion Neurons. PLoS One, 10(5), e0127572. 10.1371/journal.pone.0127572

142. Wei, X. H., Na, X. D., Liao, G. J., Chen, Q. Y., Cui, Y., Chen, F. Y., Li, Y. Y., Zang, Y., & Liu, X. G. (2013). The up-regulation of IL-6 in DRG and spinal dorsal horn contributes to neuropathic pain following L5 ventral root transection. Exp Neurol, 241, 159–168. 10.1016/j.expneurol.2012.12.007

143. Wermeling, D. P. (2005). Ziconotide, an intrathecally administered N-type calcium channel antagonist for the treatment of chronic pain. Pharmacotherapy, 25(8), 1084–1094. 10.1592/phco.2005.25.8.1084

144. White, D. M., & Cousins, M. J. (1998). Effect of subcutaneous administration of calcium channel blockers on nerve injury-induced hyperalgesia. Brain Res, 801(1-2), 50–58. 10.1016/s0006-8993(98)00539-3

145. Wong, H. S., Chang, C. M., Liu, X., Huang, W. C., & Chang, W. C. (2016). Characterization of cytokinome landscape for clinical responses in human cancers. Oncoimmunology, 5(11), e1214789. 10.1080/2162402X.2016.1214789

146. Woolf, C. J., Allchorne, A., Safieh-Garabedian, B., & Poole, S. (1997). Cytokines, nerve growth factor and inflammatory hyperalgesia: the contribution of tumour necrosis factor alpha. Br J Pharmacol, 121(3), 417–424. 10.1038/sj.bjp.0701148

147. Xia, R. H., Yosef, N., & Ubogu, E. E. (2010). Selective expression and cellular localization of pro-inflammatory chemokine ligand/receptor pairs in the sciatic nerves of a severe murine experimental autoimmune neuritis model of Guillain-Barre syndrome. Neuropathol Appl Neurobiol, 36(5), 388–398. 10.1111/j.1365-2990.2010.01092.x

148. Yu, L., Yang, F., Luo, H., Liu, F. Y., Han, J. S., Xing, G. G., & Wan, Y. (2008). The role of TRPV1 in different subtypes of dorsal root ganglion neurons in rat chronic inflammatory nociception induced by complete Freund’s adjuvant. Mol Pain, 4, 61. 10.1186/1744-8069-4-61

149. Zeng, X., Lu, S., Li, M., Zheng, M., Liu, T., Kang, R., Xu, L., Xu, Q., Song, Y., & Liu, C. (2022). Inflammatory Cytokine-Neutralizing Antibody Treatment Prevented Increases in Follicular Helper T Cells and Follicular Regulatory T Cells in a Mouse Model of Arthritis. J Inflamm Res, 15, 3997–4011. 10.2147/JIR.S355720

150. Zhang, J. M., & An, J. (2007). Cytokines, inflammation, and pain. Int Anesthesiol Clin, 45(2), 27–37. 10.1097/AIA.0b013e318034194e

151. Zhang, R. X., Li, A., Liu, B., Wang, L., Ren, K., Zhang, H., Berman, B. M., & Lao, L. (2008). IL-1ra alleviates inflammatory hyperalgesia through preventing phosphorylation of NMDA receptor NR-1 subunit in rats. Pain, 135(3), 232–239. 10.1016/j.pain.2007.05.023

152. Zhang, S., Li, Y., & Tu, Y. (2021). Lidocaine attenuates CFA-induced inflammatory pain in rats by regulating the MAPK/ERK/NF-kappaB signaling pathway. Exp Ther Med, 21(3), 211. 10.3892/etm.2021.9643

153. Zhang, W., Xiao, D., Mao, Q., & Xia, H. (2023). Role of neuroinflammation in neurodegeneration development. Signal Transduct Target Ther, 8(1), 267. 10.1038/s41392-023-01486-5

154. Zhou, Y. Q., Liu, Z., Liu, Z. H., Chen, S. P., Li, M., Shahveranov, A., Ye, D. W., & Tian, Y. K. (2016). Interleukin-6: an emerging regulator of pathological pain. J Neuroinflammation, 13(1), 141. 10.1186/s12974-016-0607-6

155. Zumerle, S., Cali, B., Munari, F., Angioni, R., Di Virgilio, F., Molon, B., & Viola, A. (2019). Intercellular Calcium Signaling Induced by ATP Potentiates Macrophage Phagocytosis. Cell Rep, 27(1), 1–10 e14. 10.1016/j.celrep.2019.03.011

