## Supplementary material for "Peripheral Ca_V_2.2 channels in skin regulate prolonged heat hypersensitivity during neuroinflammation": Salib_Supplementary Figures and Tables 13July2024

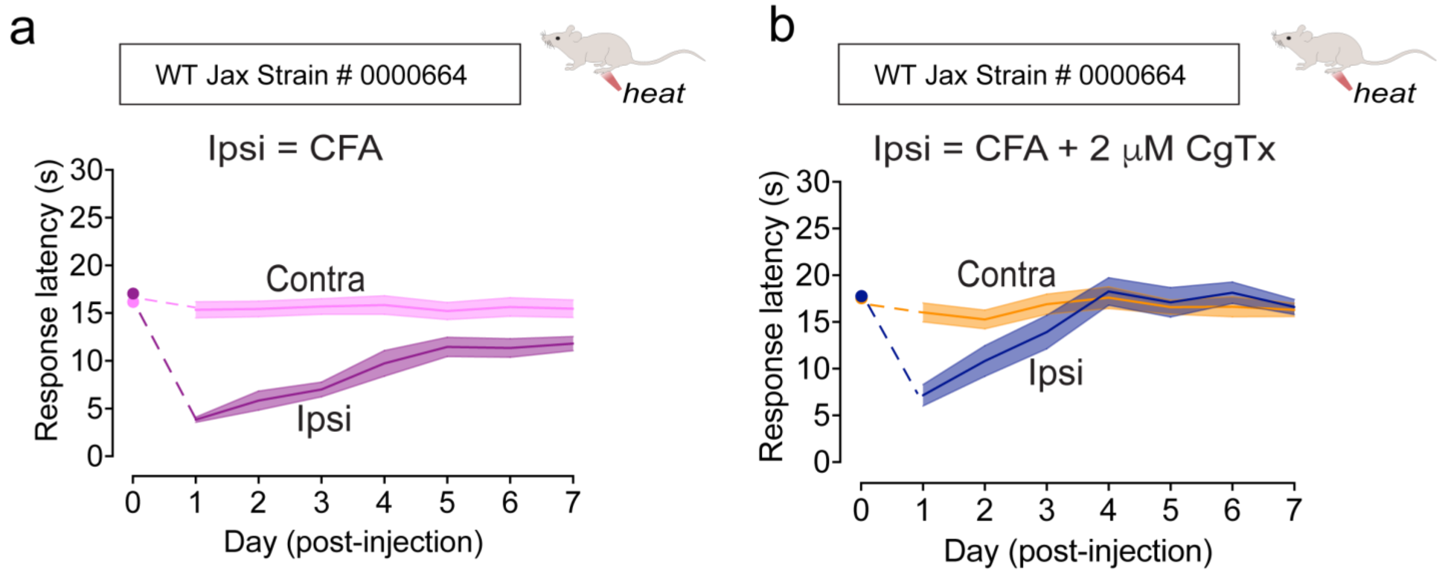


**Supplementary Figure S1** Validation of CFA-induced heat hypersensitivity inhibited by pharmacological block of peripheral $Ca_V2.2$ channels in an outbred independent mouse strain. 16–20-week-old C57BL/6J mice were ordered from Jackson Laboratories (Strain # 0000664) and were acclimated at least one week prior to behavioral testing. Under blinded conditions, mice were injected with CFA alone (n=13), or CFA + 2 μM ω-CgTx MVIIA (n=13) as described previously (see in methods and figure 2. Solid line represents the mean response latency, the shaded area represents standard error.

**Supplementary Figure S2**

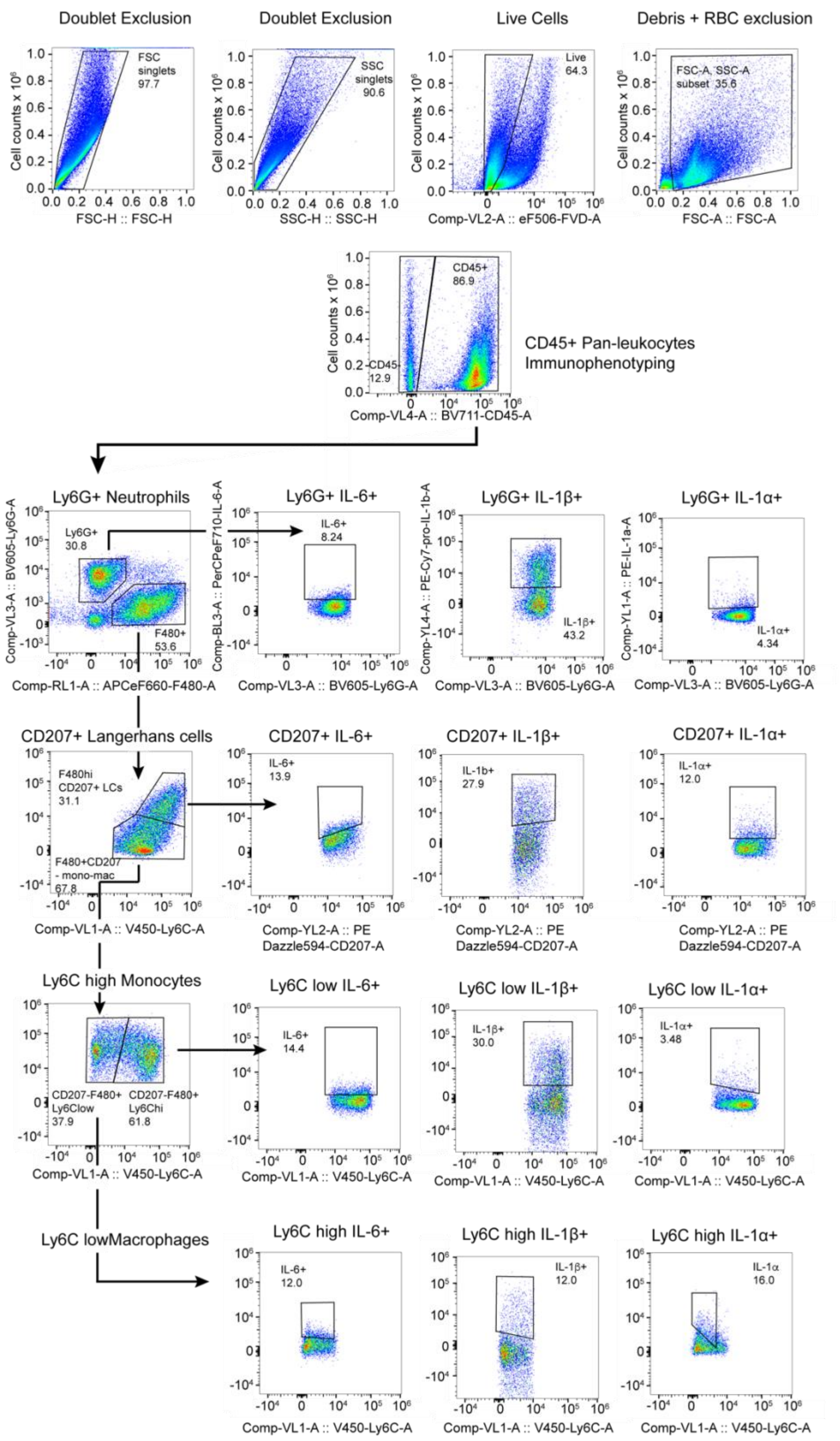

**Supplementary Figure S2** Gating strategy for hind paw deep punch biopsy immunophenotyping flow cytometry analyses (shown in Fig. 2) Analyses were performed using FlowJo Software v10.9.0. Housekeeping exclusion of cell doublets, dead cells, debris, and red blood cells. Cells were assigned an identity based on expression of specific cell surface markers using antibodies with distinct fluorophores. The cell surface marker CD45 was used to identify pan-leukocytes, that were further characterized into individual cell populations based on the expression of additional cell surface markers: Neutrophils express Ly6G, Langerhans cells express F4/80 and CD207, Monocytes express F4/80 and high expression of Ly6C, and Macrophages express F4/80 and low expression of Ly6C. For each leukocyte population, we analyzed intracellular cytokine levels of IL-6, IL-1$\beta$, and IL-1$\alpha$ using intracellular cytokine antibodies (See Table 1 and Methods).

### MSD SUPPLEMENT

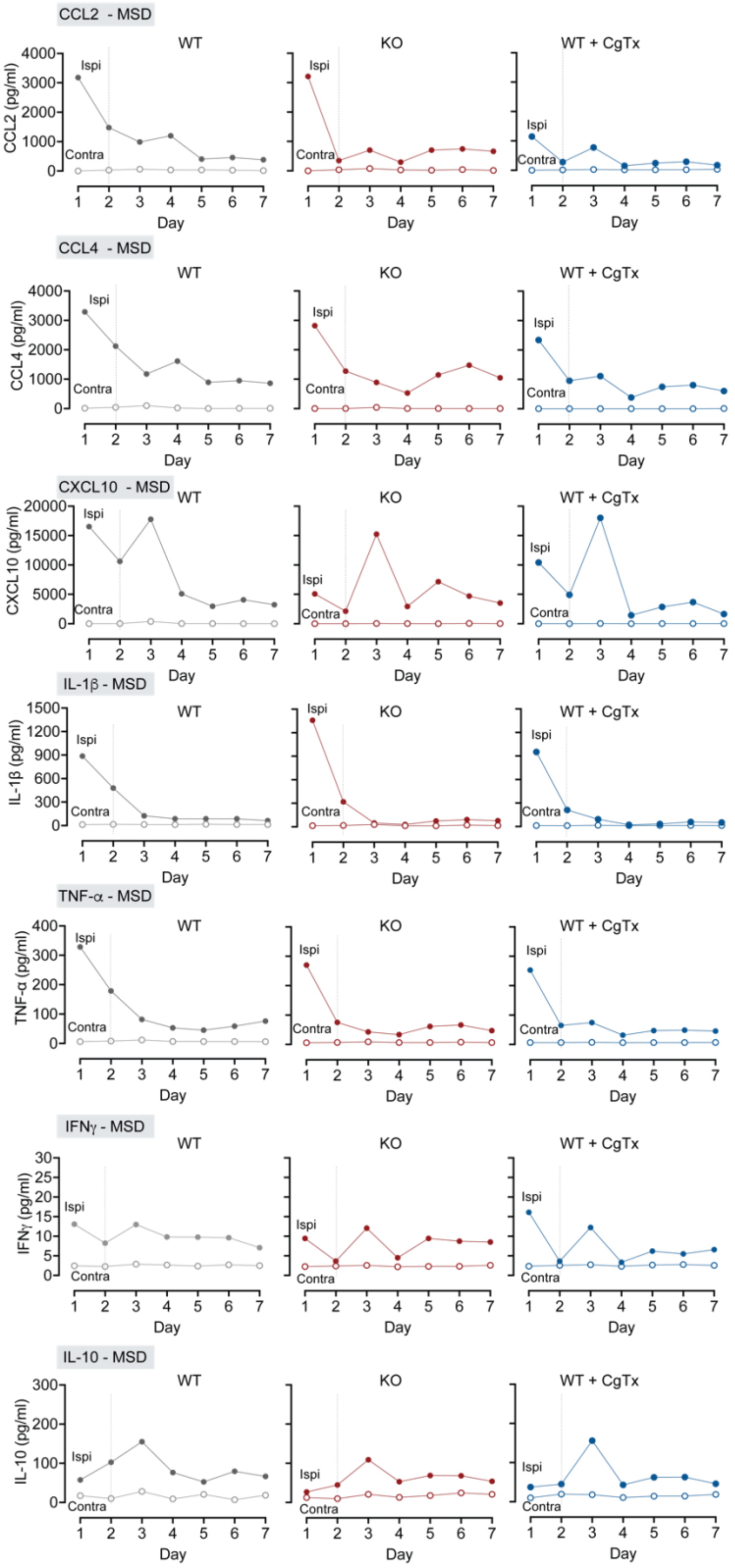

### LEGENDplex SUPPLEMENT

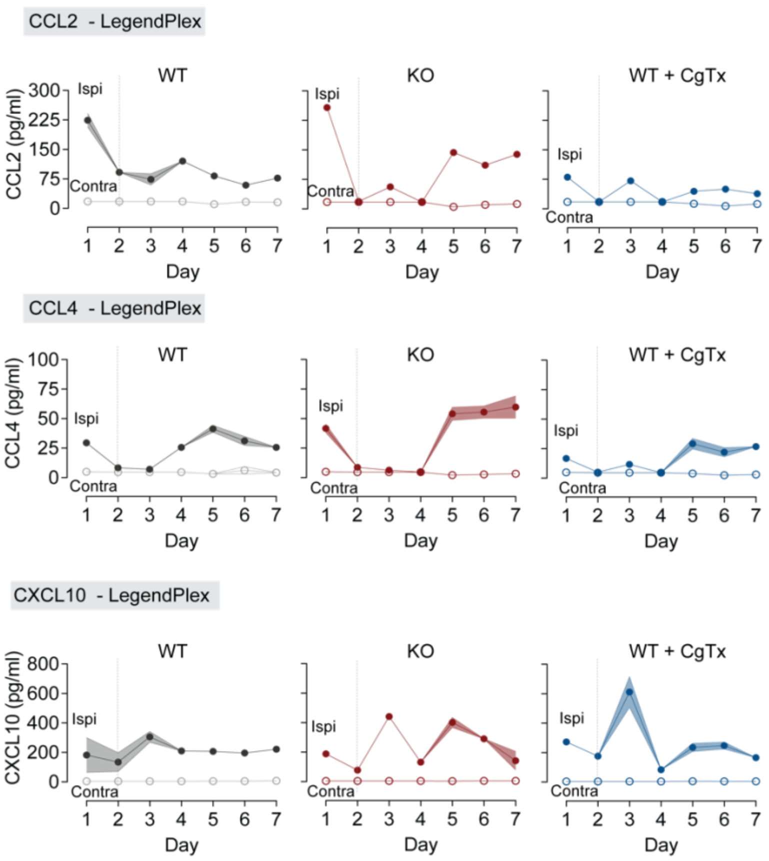


**Supplementary Figure S3** Cytokine levels in contralateral (control) and ipsilateral hind paws of mice measured daily for 1 week following 20 μL intradermal (*id*) injection of CFA in wildtype (WT, gray), CFA in $Ca_V2.2^{-/-}$ (KO, red), and CFA in WT co-injected with 2 μM ω-CgTx MVIIA (WT + CgTx, blue). N=8 mice for each condition. Cytokine detection was performed using both LEGENDplex and MSD assays. **MSD SUPPLEMENT:** Electrochemiluminescence multiplex spot-based immunoassay (MSD R-Plex, U-plex) validation unveiled four additional cytokines: IL-1β, TNF-α, IFNγ, and IL-10 in paw lavage fluid. Samples were measured in technical replicates. In addition to the five cytokines IL-6, IL-1α**,** CCL2, CCL4, and CXCL10, the MSD platform detected IL-1β, TNF-α IFNγ, and IL-10 in paw lavage fluid. **LEGENDplex SUPPLEMENT**: Samples were measured in technical triplicates. Mean ±

SE levels of CCL2 in ipsilateral paws on day 1 WT = 224 ± 17 pg/ml; KO = 258 ± 7.7 pg/ml; WT + CgTx MVIIA = 80 ± 4.6 pg/ml; day 2 WT = 92 ± 4.1 pg/ml; KO = 19 ± 0.9 pg/ml; WT + CgTx MVIIA = 17.5 ± 0.1 pg/ml; day 3: WT = 74 ± 14.2 pg/ml; KO = 56 ± 3.9 pg/ml; WT + CgTx MVIIA = 71.1 ± 5.1 pg/ml. p(WT/ KO/ CgTx MVIIA) | time interaction p<0.0001. Analysis of variance of ipsilateral paw measured using two-way ANOVA. Mean ± SE levels of CCL4 in ipsilateral paws on day 1 WT = 29.4 ± 2.2 pg/ml; KO = 41.6 ± 3.5 pg/ml; WT + CgTx MVIIA = 16 ± 1.8 pg/ml; day 2 WT = 8.2 ± 0.4 pg/ml; KO = 8.7 ± 0.3 pg/ml; WT + CgTx MVIIA = 5.1 ± 0.5 pg/ml; day 3: WT = 7.0 ± 1.3 pg/ml; KO = 6.3 ± 0.3 pg/ml; WT + CgTx MVIIA = 11.6 ± 1.8 pg/ml. p(WT/ KO/ CgTx MVIIA) | time interaction p<0.0001. Analysis of variance of ipsilateral paw measured using two-way ANOVA. **f.** Mean ± SE levels of CXCL10 in ipsilateral paws on day 1 WT = 182.3 ± 117.4 pg/ml; KO = 188.6 ± 9.3 pg/ml; WT + CgTx MVIIA = 273.1 ± 13.2 pg/ml; day 2 WT = 134.5 ± 63.5 pg/ml; KO = 77.7 ± 1.3 pg/ml; WT + CgTx MVIIA = 176.3 ± 20.9 pg/ml; day 3: WT = 305 ± 36.8 pg/ml; KO = 441.1 ± 10.6 pg/ml; WT + CgTx MVIIA = 611.4 ± 105 pg/ml. p(WT/ KO/ CgTx MVIIA) | time interaction p=0.0005. Analysis of variance of ipsilateral paw measured using two-way ANOVA.

| Cytokine | References |
|---|---|
| IL-1α | (Dinarello, 2011); (Dinarello & van der Meer, 2013); (Honore et al., 2006); (Sommer et al., 1999); (Cavalli et al., 2021) |
| IL-1β | (Dinarello, 2011); (Dinarello & van der Meer, 2013); (Honore et al., 2006); (Sommer et al., 1999); (Ahn et al., 2005; Binshtok et al., 2008; Cumberbatch et al., 2002; Ebbinghaus et al., 2012; Macleod et al., 2021; Safieh-Garabedian et al., 1995; Stemkowski & Smith, 2012) |
| IL-4 | (Huntula et al., 2019); (Junttila, 2018); (Sandy-Hindmarch et al., 2022); (Zumerle et al., 2019) |
| IL-6 | (Scholz & Woolf, 2007); (Zhou et al., 2016); (Andratsch et al., 2009; DeLeo et al., 1996; Kang et al., 2019; Latourte et al., 2017; Lin et al., 2017; Malsch et al., 2014; McLoughlin et al., 2004; Melemedjian et al., 2014; Wang et al., 2009; Wei et al., 2013) |
| IL-10 | (Chen et al., 2010); (Scholz & Woolf, 2007); (Huntula et al., 2019) |
| TNF-α | (Homma et al., 2002); (Woolf et al., 1997); (Cunin et al., 2011; Ebersberger, 2018; Liu et al., 2007; Lonnemann et al., 1989; Scholz & Woolf, 2007; Shamash et al., 2002) |
| LIF | (Banner et al., 1998); (Chen et al., 2021); (Scholz & Woolf, 2007) |
| CXCL10 | (Chen et al., 2019); (Xia et al., 2010); (Hua et al., 2021) |
| CCL2 | (Van Steenwinckel et al., 2015); (Dansereau et al., 2021); (Dansereau et al., 2021; Menetski et al., 2007; Schober, 2008; Van Steenwinckel et al., 2015) |
| CCL4 | (Aguirre et al., 2020); (Green et al., 2019), (Jiang et al., 2020) |
| IFNγ | (Adefegha et al., 2020);;(Sonekatsu et al., 2016); (Scholz & Woolf, 2007) |
| MDC | (Oh et al., 2001); (Silva et al., 2022) |
| IL-33 | (Huang et al., 2020), (Huang et al., 2021) |

**Supplementary Table 1:** Literature implicating cytokines in maladaptive prolonged inflammation.

| LEGENDplex | MSD |
|---|---|
| IL-1β | IL-1β |
| IFNγ | IFNγ |
| TNF-α | TNF-α |
| CCL4 | CCL4 |
| CCL2 | CCL2 |
| CXCL10 | CXCL10 |
| IL-6 | IL-6 |
| IL-1α | IL-1α |
| IL-10 | IL-10 |
| IL-23 | IL-23 |
| **LIF** | **MDC** |
| **IL-4** | **IL-33** |

**Supplementary Table 2:** 12-plex for LEGENDplex and MSD immunoassays. After initial pilot screens, two custom panels were developed to assess 12 cytokines across two platforms, 10 of the same cytokines were assessed on both platforms, and 2 were unique to each custom panel (bold lavender- LEGENDplex, bold blue- MSD).
